# Vps68 cooperates with ESCRT-III in intraluminal vesicle formation

**DOI:** 10.1101/2022.01.03.474785

**Authors:** Sören Alsleben, Ralf Kölling

## Abstract

The endosomal sorting complex required for transport (ESCRT)-III mediates budding and abscission of intraluminal vesicles (ILVs) into multivesicular endosomes. To further define the role of the ESCRT-III associated protein Mos10/Vps60 in ILV formation, we screened for new interaction partners by SILAC/MS. Here, we focused on the newly identified interaction partner Vps68. Our data suggest that Vps68 cooperates with ESCRT-III in ILV formation. The deletion of *VPS68* caused a sorting defect similar to the *SNF7* deletion, when the cargo load was high. The composition of ESCRT-III was altered, the level of core components was higher and the level of associated proteins was lower in the deletion strain. This suggests that a shift occurs from an active complex to a disassembly competent complex and that this shift is blocked in the *Δvps68* strain. We present evidence that during this shift Snf7 is replaced by Mos10. Vps68 has an unusual membrane topology. Two of its potential membrane helices are amphipathic helices localized to the luminal side of the endosomal membrane. Based on this membrane topology we propose that Vps68 and ESCRT-III cooperate in the abscission step by weakening the luminal and cytosolic leaflets of the bilayer at the abscission site.

## Introduction

Endosomal sorting complex required for transport (ESCRT)-III proteins are membrane remodeling factors that are involved in the deformation and abscission of membranes. ESCRT-III participates in a large number of membrane-related cellular processes (Gatta & Carlton, 2019; Hurley, 2015; McCullough *et al*, 2018; Vietri *et al*, 2020). Initially, ESCRT-III proteins were identified in yeast as factors involved in the formation of intraluminal vesicles (ILVs) at late endosomes (Babst *et al*, 2002). But now it is clear that ESCRT-III proteins are an inherent part of all eukaryotic cells. ESCRT-III proteins are of ancient origin, they were already present in the last common eukaryotic ancestor (LCEA) (Leung *et al*, 2008). Recent reports indicate that the ESCRT-III function may be even more general and widespread than initially thought. Cryo-EM studies show that the bacterial phage shock protein PspA and the plant chloroplast protein Vipp1 share a common evolutionary origin with ESCRT-III proteins (Junglas *et al*, 2021; Liu *et al*, 2021). Most ESCRT-III proteins have the propensity to form filaments. The structure of the ESCRT-III assemblies has been meticulously studied *in vitro* (Banjade *et al*, 2019; Bertin *et al*, 2020; Chiaruttini *et al*, 2015; Lee *et al*, 2015; Maity *et al*, 2019; Tang *et al*, 2015), but how ESCRT-III looks like *in vivo* is still unclear. If ESCRT-III forms structures similar to PspA and Vipp1 the complex may be less extensive as suggested by the previous *in vitro* studies.

In yeast, the role of ESCRT-III in the formation of intraluminal vesicles (ILVs) has been most thoroughly studied. It is thought that the recruitment of ESCRT-III to the endosomal membrane is the final step in a cascade of reactions involving the upstream complexes ESCRT-0, -I and -II (Teis *et al*, 2008). The common view is that ESCRT-III itself is assembled in a stepwise manner. First, Vps20 is recruited by binding to the ESCRT-II subunit Vps25, then Snf7 enters and is induced to polymerize. Snf7 polymerization is limited by Vps2 and Vps24, which finally initiate disassembly of the complex by the AAA-ATPase Vps4. This view has been challenged recently (Adell *et al*, 2017; Mierzwa *et al*, 2017). According to these newer findings, there seems to be a continuous, stochastic exchange of ESCRT-III proteins and Vps4 with sites of ILV formation. Thus, ESCRT-III appears to be far more dynamic than originally thought.

The ESCRT-III protein family consists of eight members in yeast and twelve members in mammalian cells. With respect to ILV formation, the ESCRT-III proteins are conventionally divided into two groups, the so-called core subunits Snf7, Vps2, Vps20 and Vps24 (ESCRT-III proper) and the ESCRT-III associated proteins Did2, Ist1 and Mos10/Vps60 (Azmi *et al*, 2008; Dimaano *et al*, 2008; Rue *et al*, 2008). The eighth member of the family Chm7 is not involved in ILV formation, but rather plays a separated role at the nuclear membrane (Bauer *et al*, 2015; Thaller *et al*, 2019). Our recent data suggest that the distinction between core components and associated ESCRT-III proteins may not be justified (Heinzle *et al*, 2019). We think that Did2 and Mos10 are an integral part of ESCRT-III. This view is supported by another study, which investigated the contribution of Did2 and Ist1 to ESCRT-III function (Pfitzner *et al*, 2020).

There are conflicting reports as to the role of the so-called ESCRT-III associated proteins. In our initial identification and characterization of Mos10/Vps60, the *MOS10* deletion mutant had a sorting defect indistinguishable from the deletion of other ESCRT-III core subunits (Kranz *et al*, 2001). In another report, the *MOS10* deletion only showed a sorting effect in combination with deletions of other ESCRT-III associated factors (Rue *et al.*, 2008). Recently, at least a partial sorting defect of the *MOS10* deletion has been reported in several studies (Banjade *et al*, 2021; Nickerson *et al*, 2010).

The impact of ESCRT-III deletions on the morphology of multivesicular endosomes was studied by EM tomography (Nickerson *et al*., 2010; Nickerson *et al*, 2006). From these studies, a clear distinction could be made between core subunits and associated proteins. Deletion of core subunits completely prevented ILV formation and led to the accumulation of stacked membrane structures next to the vacuole (the so-called class E compartments) (Raymond *et al*, 1992). In contrast, deletion of associated factors gave rise to multivesicular endosomes with a vesicular/tubular morphology (VTEs) (Nickerson *et al*., 2010). Thus, the associated factors seem to act after the core components. Evidence has been presented that the associated proteins Did2 and Mos10 are involved in the disassembly of ESCRT-III (Azmi *et al.,* 2008; Dimaano *et al*., 2008; Rue *et al*., 2008). Our interpretation of these findings is that ESCRT-III can exist in different functional states with different compositions. At the beginning of the functional cycle ESCRT-III is actively involved in ILV formation and/or abscission, then after completion of ILV formation, it is converted to a disassembly competent complex. The data presented in this report are in line with this interpretation.

To learn more about the function of Mos10, we looked for new interaction partners by a SILAC/MS screen. Consistently, the ESCRT-III proteins Vps2, Vps24, Snf7 and Did2 were copurified with Mos10. This strengthens the view that Mos10 is an integral part of ESCRT-III. In addition, several other potential candidates were detected. Here, we focused on one of these interaction partners, the protein Vps68. Vps68 was first identified in a screen for vacuolar protein sorting (*vps*) mutants that mis-localize carboxypeptidase Y (Bonangelino *et al*, 2002). By genome-wide screens, it could be shown that Vps68 localizes to endosomes and that it forms a complex with Vps55 (Huh *et al*, 2003; Schluter *et al*, 2008).

Here we show that Vps68 physically interacts with ESCRT-III and present evidence that it cooperates with ESCRT-III in ILV formation.

## Results

### Screen for Mos10 interacting proteins

Although, the ESCRT-III system has been intensively studied, there are only relatively few reports about the role of the human ESCRT-III protein CHMP5 or its yeast counterpart Mos10/Vps60. To learn more about the function of Mos10, we looked for novel Mos10 interaction partners. To this end, Mos10 was purified from yeast cell extracts by affinity chromatography and the co-purified proteins were then identified by mass spectrometry. Purifications were performed with two different tags fused to the Mos10 C-terminus, a 6His-tag and a superfolder-GFP (sfGFP)-tag. Mos10-6His was functional (Fig. S6-2), while Mos10-sfGFP gave rise to a block in endocytic trafficking (Fig. S6-3). To filter out unspecific binding, a SILAC (stable isotope labeling with amino acids in cell culture) approach was used. The culture expressing tagged Mos10 was grown in medium containing arginine and lysine labeled with heavy isotopes, while the control culture expressing native Mos10 was grown with arginine and lysine with normal, light isotopes. Among the proteins identified by mass spectrometry, we looked for proteins with a high heavy to light (H/L) ratio. The results of three different purifications (two Ni-NTA purifications with the 6His-tag, one with anti-GFP antibodies and the sfGFP-tag) are summarized in Tab. S1. We detected 51 proteins that were at least 2-fold enriched for the heavy isotopes. Two of these proteins (Bag7, Ykr075c) are naturally occurring poly-histidine proteins and are thus most likely unspecific contaminants. The list of potential interaction partners was checked against the CRAPome database (Mellacheruvu *et al*, 2013) to filter out common contaminants in AP-MS experiments. Proteins detected at least 5-times in 17 experiments were dismissed as unspecific. The remaining 37 proteins are candidates for further study. The proteins with the highest H/L ratio, which were consistently co-purified in all three experiments, were the ESCRT-III proteins Vps2, Snf7, Vps24 and Did2. This clearly demonstrates that Mos10 is a bona fide member of ESCRT-III, a fact that is still not generally acknowledged. From the list of the remaining proteins, Vps68 was selected for further study.

### Vps68 binds to ESCRT-III

The binding of Vps68 to Mos10 was verified by a co-immunoprecipitation experiment. For detection, Vps68 was tagged with sfGFP at its N-terminus. N-terminally tagged Vps68 proved to be functional, while C-terminally tagged Vps68 was compromised in its function (Fig. S3). Vps68 was immunoprecipitated from cell extracts with anti-GFP antibodies and examined for co-immunoprecipitation of the ESCRT-III proteins Mos10, Snf7 and Vps2 (Fig. 1). In the otherwise wildtype background, all three ESCRT-III proteins could be co-immunoprecipitated by the GFP antibodies. This confirms our MS identification of Vps68 as an interaction partner of Mos10.

**Figure 1.**
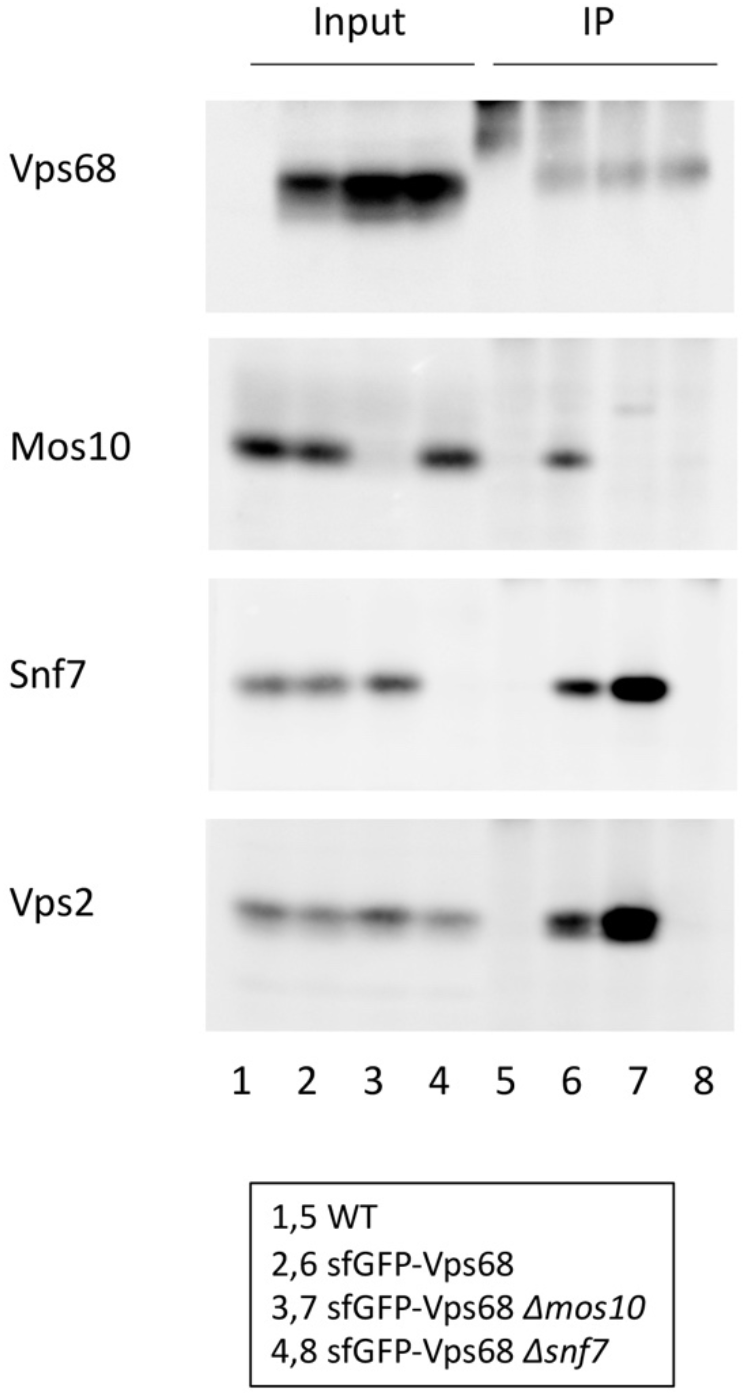
Vps68 binds to ESCRT-III. sfGFP-Vps68 was immunoprecipitated from cell extracts with anti-GFP antibodies. The immunoprecipitates were examined for the presence of ESCRT-III proteins with specific antibodies. Lanes 1-4: Input, lanes 5-8: IP. Lanes 1,5: RKY1558 (WT), lanes 2,6: RKY3285 (*sfGFP-VPS68*), lanes 3,7: RKY3394 (*sfGFP-VPS68 Δmos10*), lanes 4,8: RKY3412 (*sfGFP-VPS68 Δsnf7*)

We were then interested to know, whether Vps68 is recruited to ESCRT-III by direct binding to Mos10 or whether the interaction is mediated by some other subunit of ESCRT-III. To this end, the co-immunoprecipitation experiment was repeated with a *Δmos10* and a *Δsnf7* strain carrying the integrated *sfGFP-VPS68* cassette. As can be seen in Fig. 1, Snf7 and Vps2 could still be precipitated by Vps68 in the *Δmos10* strain. Thus, binding of Vps68 to ESCRT-III does not appear to be mediated by Mos10. In the *Δsnf7* strain, no co-precipitation of any ESCRT-III subunit could be observed. This is in line with the observation that Snf7 is absolutely essential for ESCRT-III formation (Heinzle *et al*., 2019), i.e. in *Δsnf7* there is no ESCRT-III that could be precipitated by Vps68. The amount of Snf7 and Vps2 precipitated by Vps68 was twice as high in the *Δmos10* strain compared to wildtype. Thus, ESCRT-III complexes accumulate in the cell, when Mos10 is missing. This is consistent with a role of Mos10 in disassembly of ESCRT-III.

This notion was explored further by sucrose density gradient fractionation of cell extracts from a wildtype and from a *Δmos10* strain (Fig. 2, Fig. S2). The fractionation profile of Snf7 was compared to the profiles of two marker proteins, Pep12 as a marker for endosomes and ALP as a marker for the vacuole. With wildtype extracts, Pep12 had a peak in fraction 7 and ALP had a peak in fraction 8. In the *Δmos10* strain, the Pep12 peak was shifted towards fraction 8, while the ALP peak was unchanged. With wildtype extracts, most of Snf7 was found at the top of the gradient, where the soluble proteins are localized. A small part migrated into the gradient and formed a small peak in fraction 7, consistent with an endosomal localization of Snf7. With *Δmos10* extracts, the membrane associated fraction of Snf7 was markedly higher, with a concomitant reduction of the soluble fraction. This agrees with our co-IP results and shows that Mos10 is involved in the disassembly of ESCRT-III.

**Figure 2.**
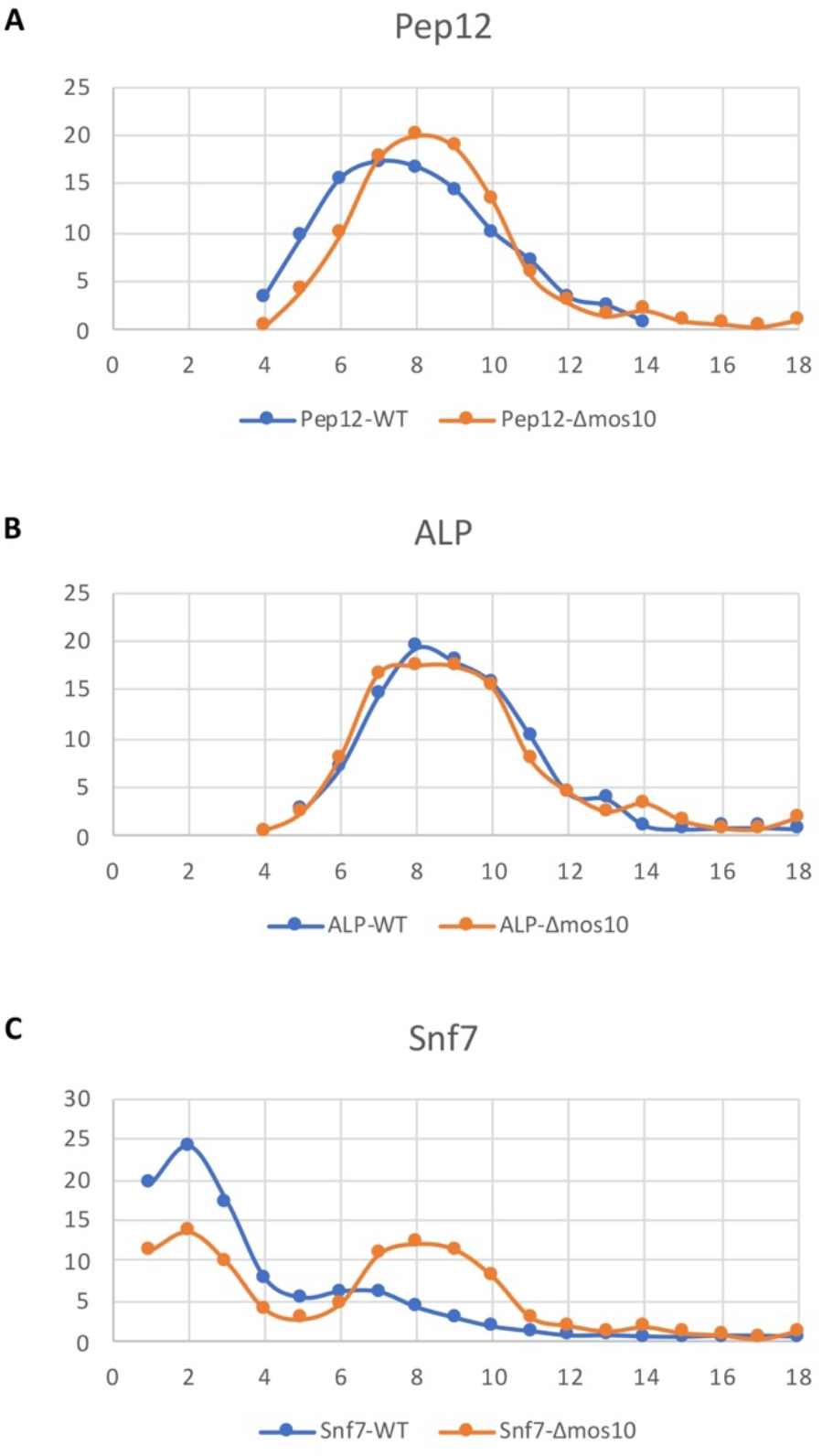
Membrane accumulation of Snf7 in a *MOS10* deletion mutant. Yeast cell extracts of the wildtype RKY1558 (blue) and the *Δmos10* strain RKY2909 (orange) were fractionated by sucrose density gradient fractionation and analyzed by western blotting. (A) Pep12, (B) Alkaline phosphatase (ALP), (C) Snf7. The western blot signals (Fig. S2) were quantified with ImageJ. The percentage of total protein per fraction is given. Left: low density, right: high density.

### Vps68 is involved in the sorting of the endocytic cargo protein Ste6

Vps68 has been identified previously as a protein involved in the trafficking of proteins to the yeast vacuole (Bonangelino *et al*., 2002; Huh *et al*., 2003; Schluter *et al*., 2008). To further explore its role in endocytic trafficking and to gain information about how Vps68 intersects with ESCRT-III function, the effect of *VPS68* deletion on the endocytic cargo protein Ste6 was examined. The ABC-transporter Ste6 is a very short-lived protein (Kölling & Hollenberg, 1994). After transport to the plasma membrane, it is internalized by endocytosis and moves to the vacuole for degradation. To see, if Vps68 is required for Ste6 transport to the vacuole, the Ste6 half-life was determined by a Gal-depletion experiment in a wildtype stain and in a *VPS68* deletion (Fig. 3). Ste6 was expressed from a *GAL1* promoter inserted in front of the chromosomal *STE6* gene. Cells were grown in galactose to induce expression from the *GAL1* promoter and were then shifted to glucose containing medium, to repress transcription. In the wildtype strain, Ste6 was degraded with a half-life of 12 min. In the *Δvps68* strain, there was a slight delay in the degradation of Ste6 after shift to glucose medium and then Ste6 was degraded with a half-life of 29 min. Thus, Ste6 is moderately stabilized in the *VPS68* deletion. Similar results had been reported for the a-factor receptor Ste3 (Schluter *et al*., 2008).

**Figure 3.**
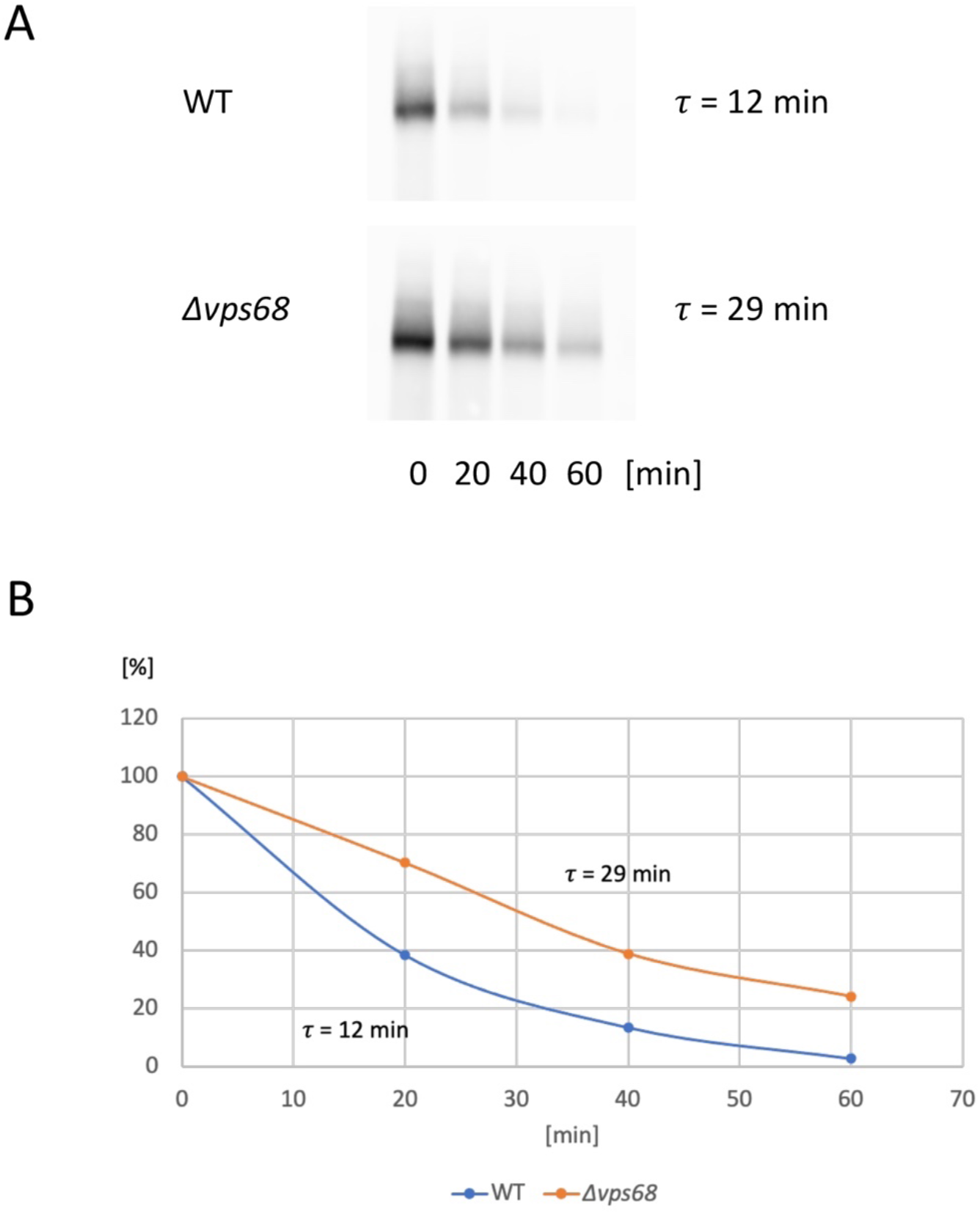
*VPS68* deletion stabilizes Ste6. Gal-depletion experiment. (A) Yeast cells were grown on 2 % galactose medium and shifted to 2 % glucose medium at t_0_. Cell extracts from equal culture volumes were prepared at the times indicated and examined for Ste6 by western blotting. (B) The western blot signals were quantified by ImageJ (t_0_=100 %). Ste6 was expressed from the *GAL1* promoter. Strains: RKY3319 (wildtype, blue), RKY3320 (*Δvps68*, orange).

To determine the site of action of Vps68 in the endocytic pathway an epistasis analysis with established markers was performed. The localization of Ste6-GFP expressed from a multicopy plasmid in different single and double mutant strains was examined by fluorescence microscopy (Fig. 4). In the wildtype, Ste6-GFP stained the lumen of the vacuole and occasionally some endosomal dots. In the class D mutant *Δvps21*, with a defect in the fusion of endocytic vesicles with the endosome (Bowers & Stevens, 2005), Ste6-GFP displays enhanced recycling to the yeast cell surface (Krsmanović *et al*, 2005). In this mutant, a number of smaller vesicles, concentrated at the bud-neck and some staining of the bud surface, could be seen. No staining of the vacuolar lumen was visible. The *Δsnf7* mutant displayed the classical class E phenotype with a large patch at the vacuole and staining of the vacuolar membrane (Raymond *et al*., 1992). Curiously, the phenotype of the *VPS68* deletion was dependent on the expression level of Ste6-GFP. The copy number of the 2μ-vector used for expression of Ste6-GFP varies considerably, thus some cells have a very high expression level and other cells express Ste6-GFP only at low levels. With low level expression, the staining resembled wildtype, with a somewhat more pronounced staining of endosomal dots. In contrast, cells with high expression levels showed a brightly staining patch at the vacuole with a faint staining of the vacuolar membrane, similar to a class E staining.

**Figure 4.**
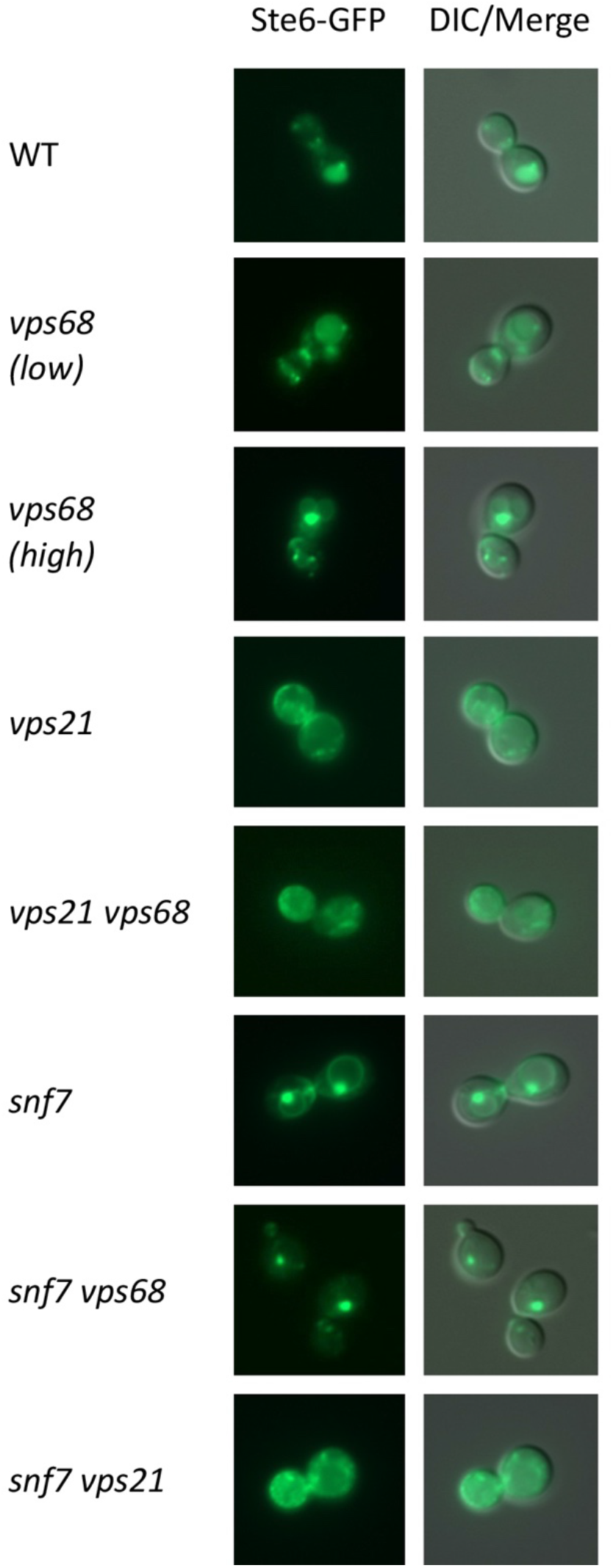
Influence of *VPS68* deletion on Ste6-GFP localization. Different yeast strains were transformed with the Ste6-GFP plasmid pRK599. The localization of Ste6-GFP was examined by fluorescence microscopy. Strains (from top to bottom): RKY1558 (WT), RKY3222 (*Δvps68*, low expression level of Ste6-GFP), RKY3222 (*Δvps68*, high expression level of Ste6-GFP), RKY1920 (*Δvps21*), RKY3307 (*Δvps21 Δvps68*), RKY2790 (*Δsnf7*), RKY3300 (*Δsnf7 Δvps68*), RKY3376 (*Δsnf7Δvps21*). Left panels: Ste6-GFP fluorescence, right panels: merged picture of differential interference contrast (DIC) image and fluorescence image.

For the epistasis analysis, the double mutants were examined. The *Δvps21 Δvps68* double mutant clearly looked like the *Δvps21* single mutant, thus *Δvps21* is epistatic over the *Δvps68* mutant, i.e. Vps21 acts before Vps68. A similar result was obtained for the *Δsnf7 Δvps21* double mutant, which resembled the *Δvps21* single mutant. When *Δsnf7* and *Δvps68* were combined, a class E-like staining was observed with the notable difference that the staining of the vacuolar membrane was missing. This synthetic phenotype indicates that Vps68 and Snf7 act at the same step.

We also examined the localization of sfGFP-Vps68 (Fig. S4). In agreement with a previous report (Schluter *et al*., 2008), we detected a handful of distinct dots, which presumably correspond to endosomes. In a *Δsnf7* background Vps68 was trapped in class E structures. Thus, it is very likely that Vps68 acts at the level of the endosome, as reported previously.

### The *VPS68* deletion alters ESCRT-III composition

To further explore the relationship between Vps68 and ESCRT-III, we looked for differences in the composition of ESCRT-III in *Δvps68* compared to wildtype. ESCRT-III was immunoprecipitated from cell extracts with antibodies against Did2, Ist1, Mos10, Snf7, Vps2 and Vps24 (Chm7 was omitted, because it is not part of the endosomal complex and Vps20 was omitted due to low signal intensity). The immunoprecipitates were then analyzed for coimmunoprecipitation of the other ESCRT-III proteins. The Co-IP signals were quantified and the wildtype Co-IP efficiencies were subtracted from the Co-IP efficiencies of the *Δvps68* strain (Fig. 5, blots in Fig. S5). As can be seen in Fig.5, a striking division among the different ESCRT-III proteins was observed. While the complexes precipitated from *Δvps68* contained less Did2, Ist1 and Mos10 compared to wildtype, they contained more Snf7, Vps2 and Vps24. Snf7, Vps2 and Vps24 are the core ESCRT-III components, which are thought to be actively involved in ILV formation and abscission. Did2, Ist1 and Mos10 are the ESCRT-III associated proteins, which are thought to be involved in disassembly of ESCRT-III (Azmi *et al*., 2008; Dimaano *et al*., 2008; Rue *et al*., 2008). Our results suggest that the *VPS68* deletion interferes with the transition from an ILV-forming ESCRT-III complex to a disassembly competent complex.

**Figure 5.**
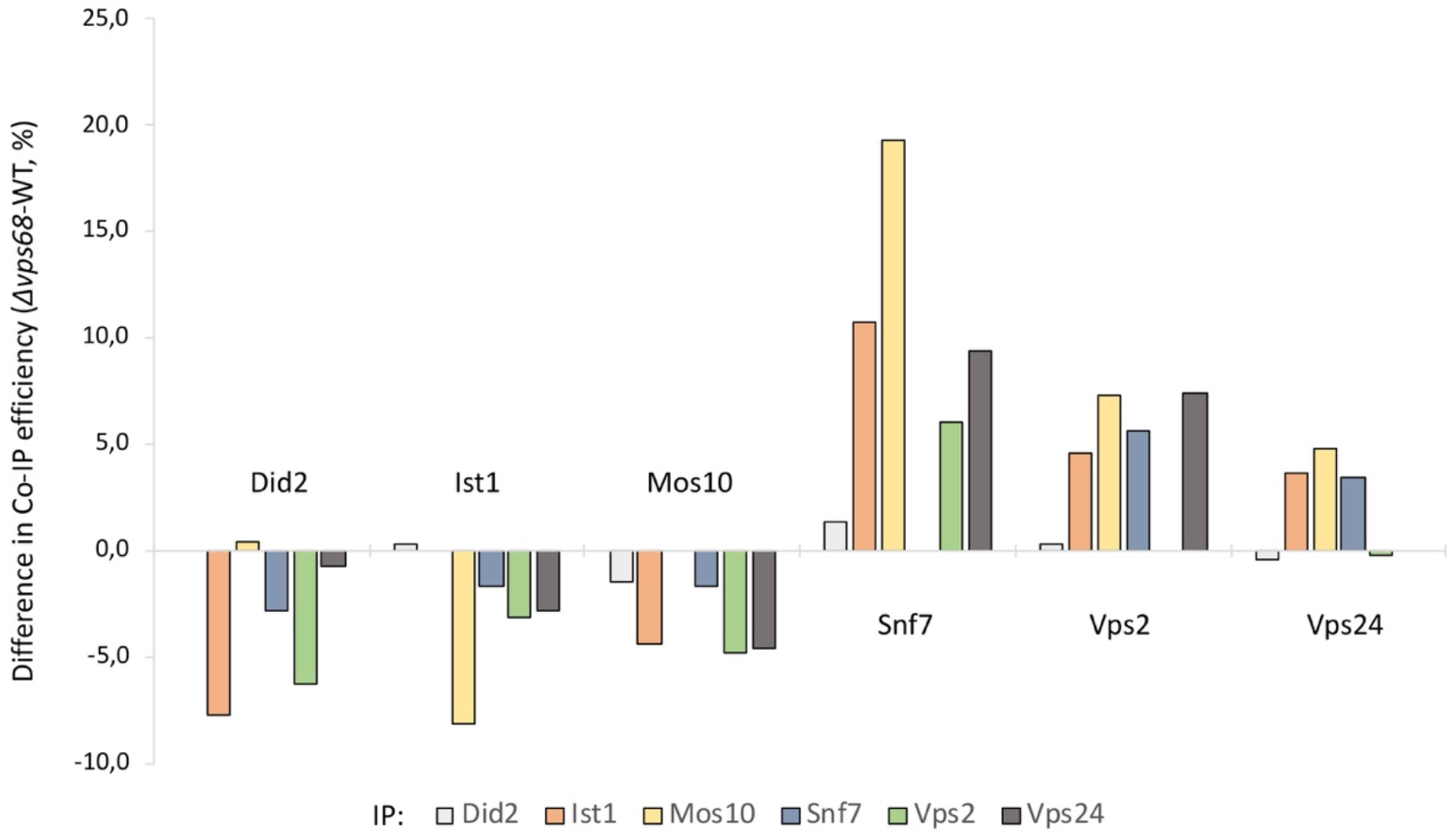
Altered ESCRT-III composition in *Δvps68*. Did2, Ist1, Mos10, Snf7, Vps2 and Vps24 were immunoprecipitated from cell extracts of RKY1558 (wildtype) and RKY3222 (*Δvps68*). The immunoprecipitates were examined for co-immunoprecipitation of other ESCRT-III proteins by western blotting. The co-IP signals (Figs. S5) were quantified by ImageJ. The percentage of protein that could be co-immunoprecipitated was calculated (= Co-IP efficiency). Co-IP efficiencies were normalized to the primary IP efficiency of the bait protein. The co-IP efficiencies of the wildtype strain were subtracted from the co-IP efficiencies of the *Δvps68* strain. The co-immunoprecipitated proteins are indicated on top or below the columns. The color code of the proteins precipitated in the primary IP is given at the bottom of the diagram (marked with IP). Average of two independent experiments.

### Effect of *VPS68* deletion on the localization of ESCRT-III proteins

Vps68 could act as a scaffold or adapter for ESCRT-III membrane association. Therefore, we examined the effect of the *VPS68* deletion on membrane association of ESCRT-III proteins by a flotation experiment (Fig. S6-1). Only minor differences in the membrane associated fractions between *Δvps68* and wildtype were observed. In general, about 30-50 % of the ESCRT-III proteins turned out to be membrane associated. The only exception was Mos10 with a very low level of membrane association of about 10 %. This experiment shows that Vps68 is not required for membrane association of ESCRT-III. Still, it could affect the site of membrane association of ESCRT-III proteins within the cell.

To examine the intracellular localization in the *VPS68* deletion versus wildtype, all ESCRT-III proteins were C-terminally tagged with sfGFP by the insertion of a tagging cassette into the yeast genome. Since it is known that tagging of ESCRT-III proteins can lead to non-functional proteins, we tested the functionality of the tagged proteins. The strains were transformed with a single copy plasmid expressing an mCherry tagged variant of the vacuolar carboxypeptidase S (Cps1). Cps1 is transported to the lumen of the vacuole via the MVB pathway. The localization of mCherry-Cps1 was examined by fluorescence microscopy (Fig. S6-2 and Fig. S6-3). In the wildtype strain, mCherry-Cps1 exclusively stained the lumen of the vacuole. In the *Δvps24* mutant as a reference a typical class E staining was observed. The strain expressing Mos10-6His displayed a wildtype pattern, showing that the tagged protein is functional. The ESCRT-III-sfGFP fusions were compromised in their function to different degrees. While the mCherry-Cps1 staining in the *IST1-sfGFP* strain was indistinguishable from wildtype, the *SNF7-, VPS2-* and *VPS24-sfGFP* strains gave rise to a strong class E phenotype. The *DID2-* and *MOS10-sfGFP* strains showed a partial defect, with some luminal staining, but also a clearly visible vacuolar rim staining. Although, most of the tagged ESCRT-III proteins cannot support normal transport of endocytic cargo proteins to the lumen of the vacuole, they may still be useful for localization studies, because, as described below, the observed staining patterns looked pretty reasonable.

The ESCRT-III proteins could be broadly divided into two groups with respect to their sfGFP staining patterns (Fig. 6). Did2 and Ist1 looked very similar with about half a dozen distinct dots. With Snf7 and Vps24 a brightly staining cluster of vesicles and a number of less intensively staining vesicles, which tended to be concentrated at the cell cortex, were observed. Vps24 resembled Snf7 and Vps2 in the sense that it also showed a brightly staining cluster of vesicles, but in contrast to these proteins, it had less additional vesicles. These staining patterns were not affected by *VPS68* deletion. Mos10-sfGFP, in contrast, showed a unique staining pattern. The staining was concentrated around the vacuole in a patchy manner. This staining is distinct from a class E staining, since the Mos10 patches were more numerous and not so grossly enlarged. This vacuolar Mos10-sfGFP staining was completely lost in *Δvps68*. In this mutant, the pattern now resembled the Snf7 pattern.

**Figure 6.**
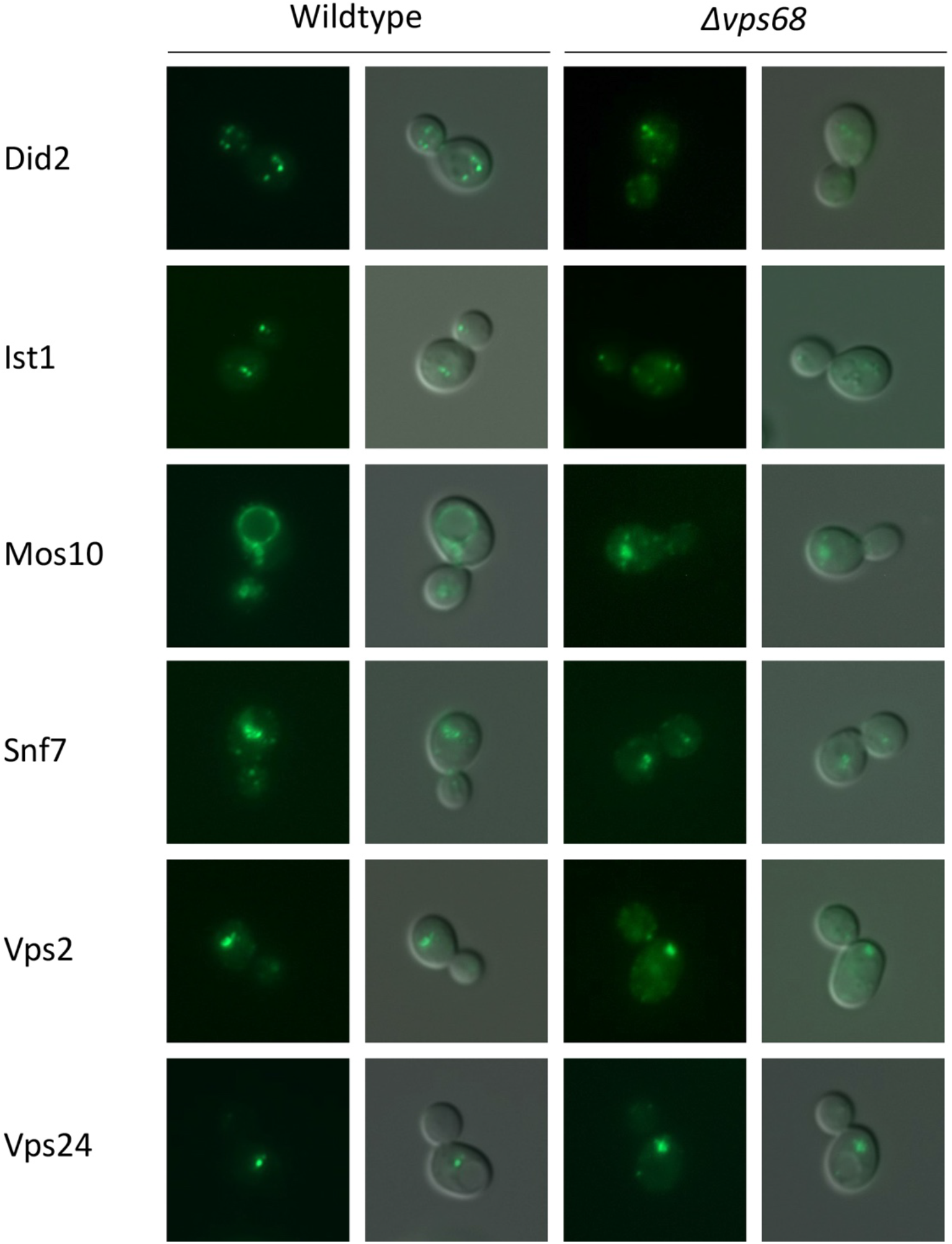
Intracellular localization of ESCRT-III-sfGFP fusions. The intracellular localization of ESCRT-III-sfGFP fusions expressed from their chromosomal loci was examined by fluorescence microscopy. Left panels: wildtype background, right panels: *Δvps68* background. For each set: GFP fluorescence and merged image (GFP+DIC). Strains from top to bottom (WT/*Δvps68*): RKY3214/3269 (Did2-sfGFP), RKY3215/3257 (Ist1-sfGFP), RKY3216/3224 (Mos10-sfGFP), RKY3217/3279 (Snf7-sfGFP), RKY3218/3271 (Vps2-sfGFP), RKY3220/3274 (Vps24-sfGFP).

To further define the dot-like structures seen with the tagged ESCRT-III proteins, we compared the localization of Snf7-mCherry with the localization of sfGFP-tagged yeast Rab proteins, which serve as markers for different organelles (Fig. S6-4). The N-terminally tagged Rab proteins were expressed from the *SNF7* promoter at their chromosomal loci. As might have been expected, Snf7-mCherry co-localized with the Rab5 homologue Vps21, a thoroughly studied endosomal marker in yeast. The brightly staining vesicular clusters were stained for both proteins. But the staining of the two proteins did not completely overlap, there were also vesicular structures that were only stained by one or the other protein (marked by arrows in Fig. S6-4). The Rab5 homologue Ypt53 closely resembled Vps21. The third Rab5 homologue Ypt52 showed an unusual staining pattern. Part of it also localized to the Snf7 clusters, but in addition, it also stained ER membranes. Thus, Ypt52 could be involved in the transport of proteins between endosomes and the ER. This intriguing finding clearly deserves further study. Other Rab proteins, like the proteins that are associated with the Golgi or post-Golgi compartments as part of the secretory pathway (Ypt6, Ypt31 and Ypt32) showed no colocalization with Snf7-mCherry. An interaction was observed between Snf7-mCherry and sfGFP-Ypt10. This protein is also considered as a Rab5 homologue and could play a role in endosome fusion to the vacuole (Langemeyer *et al*, 2020). But, so far not much is known about its function. If tagged alone, staining of the vacuolar membrane with a few associated dots was observed, reminiscent of the Mos10-sfGFP staining. However, this staining was lost, when Snf7-mCherry was present in the same strain. Apparently, Ypt10 association with endosomal and vacuolar structures requires the successful completion of the ESCRT-III function, which is prevented by Snf7 tagging.

### Mos10 accumulates in endosomal structures at the vacuole

Mos10-sfGFP shows a unique circular staining around the vacuole. Z-Stacks reveal that this staining consists of vesicular/tubular structures forming an irregular network around the vacuole (Fig. S7). These structures could result from a proliferation of the vacuolar membrane, similar to the situation of Chm7, where an accumulation inside the nucleus leads to a dramatic proliferation of the inner nuclear membrane (Thaller *et al*., 2019). Alternatively, these structures could be derived from some other cellular compartment. To distinguish between these alternatives, the localization of Mos10 was compared with the localization of marker proteins by fluorescence microscopy. In some combinations Mos10 was tagged with mCherry, in others it was tagged with sfGFP (Fig. 7). The switch of the tag was necessary, because some of the marker proteins could not be visualized by the mCherry tag due to the low fluorescence intensity. First, Mos10-sfGFP was compared with the vacuolar marker protein Vph1-mCherry. In contrast to Mos10, which displayed the usual irregular patchy staining, Vph1 showed a smooth, regular staining of the vacuolar membrane. The vacuoles were evenly stained, no concentration in patches could be seen. From this we conclude that the Mos10-sfGFP structures are not derived from the vacuolar membrane, but are instead closely associated with it. These structures do not seem to be derived from ER or Golgi membranes, since no costaining with the marker proteins Sec63 or Cog5 could be observed. With sfGFP-Vps21 a handful distinct dots were observed, which were not stained for Mos10-mCherry. In some cases, Vps21 dots were seen at the vacuolar membrane, but they did not coincide with Mos10 patches. Thus, it appears that Vps21 does not co-localize with the Mos10 structures. Next, we examined two cargo proteins of the endocytic pathway, Cps1 and Ste6, which are sorted to the vacuolar lumen via the MVB pathway. These two proteins stained the vacuolar membrane and co-localized with Mos10 patches. From this we conclude that the Mos10-sfGFP structures at the vacuolar membrane are late endosomes.

**Figure 7.**
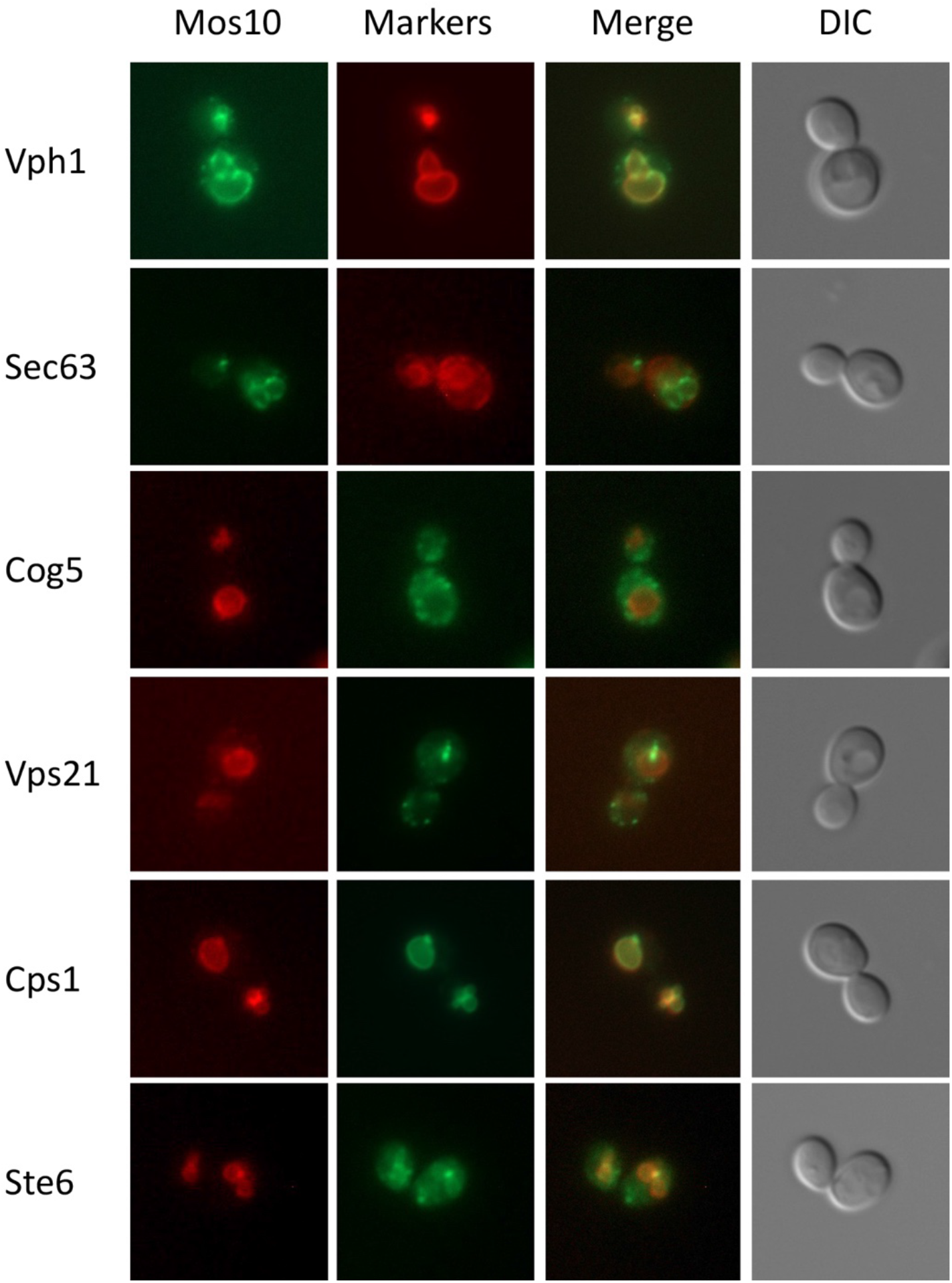
Mos10 co-localizes with endosomal cargo proteins. Yeast strains expressing a chromosomal copy of tagged versions of Mos10 and specific marker proteins were examined for co-localization by fluorescence microscopy. The proteins were either tagged with sfGFP or mCherry depending on the signal intensity of the fusion proteins involved. Panels from left to right: Mos10 fluorescence (either sfGFP or mCherry), marker protein fluorescence (either mCherry or sfGfP), merged image, DIC image. Strains from top to bottom: RKY3448 (Mos10-sfGFP, Vph1-mCherry), RKY3429 (Mos10-sfGFP, Sec63-mCherry), RKY3482 (Mos10-mCherry, Cog5-sfGFP), RKY3473 (Mos10-mCherry, sfGFP-Vps21), RKY3484 (Mos10-mCherry, Cps1-sfGFP), RKY3486 (Mos10-mCherry, Ste6-sfGFP).

### Switch from a Snf7 complex to a Mos10 complex

In a previous report, we examined the influence of ESCRT-III deletion mutations on the composition of ESCRT-III (Heinzle *et al*., 2019). We made the observation that anti-Mos10 antibodies co-immunoprecipitated most of the other ESCRT-III proteins, but not Snf7. This opened up the possibility that at a certain point in the ESCRT-III functional cycle Snf7 is replaced by Mos10, which could convert the active ESCRT-III complex into a disassembly competent complex. However, there was a caveat to this experiment. The anti-Mos10 antibodies showed a small degree of cross-reactivity towards Snf7, so we had to introduce a correction factor for this cross-reactivity. The correction factor was obtained from the anti-Mos10 IP in a *Δmos10* background. Any Snf7 signal obtained under these conditions must be derived from the direct IP of Snf7. This value was subtracted from the Snf7 co-IP signals in anti-Mos10 IPs. With this correction, no Snf7 co-IP was obtained with anti-Mos10. But there is the possibility that we over-corrected. The direct IP of Snf7 by anti-Mos10 could be lower or even non-existent in the presence of Mos10 protein.

To exclude this potential problem, we performed a pulldown experiment with Mos10-6His, which was purified from cell extracts by Ni-NTA. As a control, a Snf7-6His expressing strain was used. The proteins purified on the Ni-NTA beads were tested for the presence of ESCRT-III proteins by western blotting with specific antibodies. As can be seen in Fig. 8A, Vps2 and Vps24, but no Snf7 and barely any Did2 were co-purified with Mos10-6His. This corroborates our previous findings and shows that the performed correction was appropriate. Snf7-6His did not bring down any of the ESCRT-III proteins tested, which indicates the Snf7-6His is nonfunctional.

**Figure 8.**
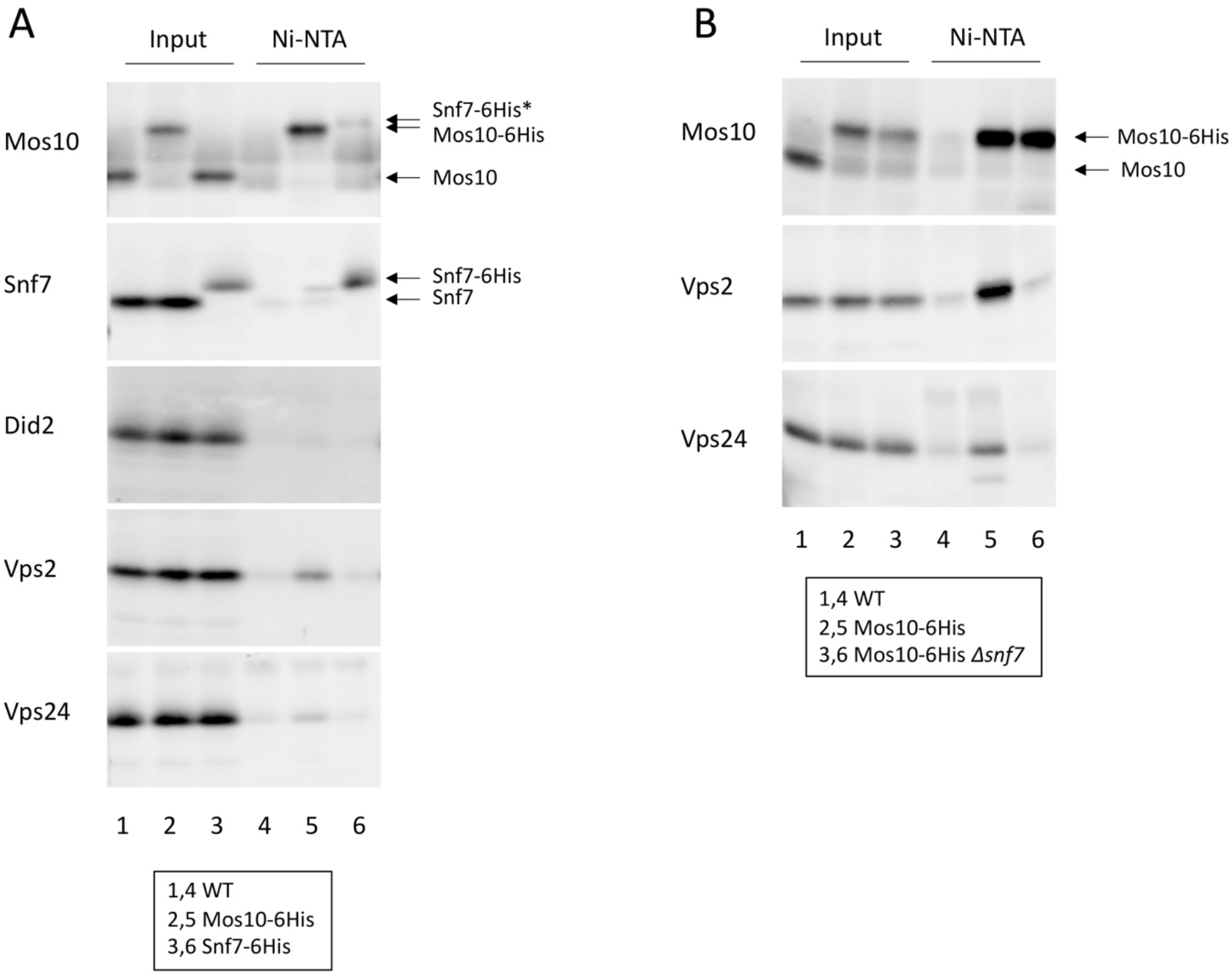
Mos10-6His and Snf7-6His pulldowns. Mos10-6His and Snf7-6His were purified from cell extracts by Ni-NTA affinity chromatography. The purified proteins were examined for co-purification of other ESCRT-III proteins by western blotting as indicated on the left side of the panels. Lanes 1-3: input, lanes 4-6: Ni-NTA pulldown. (A) The strains used were: JD52 (wildtype, lanes 1,4), RKY2889 (Mos10-6His, lanes 2,5), RKY2978 (Snf7-6His, lanes 3,6). (B) The strains used were: JD52 (wildtype, lanes 1,4), RKY2889 (Mos10-6His, lanes 2,5), RKY3438 (Mos10-6His *Δsnf7*, lanes 3,6). The position of tagged and native proteins is marked by arrows. Snf7-6His* = cross-reaction of the Mos10 antiserum with Snf7-6His.

We wondered whether the complex purified with Mos10-6His corresponded to the well-known endosomal complex involved in ILV formation or whether we purified a novel previously uncharacterized complex. To this end, we performed a pulldown experiment in a *Δsnf7* background (Fig. 8B). Again, Vps2 and Vps24 could be co-purified with Mos10-6His in the wildtype background, while no co-purification was observed in the *Δsnf7* background. This demonstrates that although Mos10-6His does not bring down any Snf7, the purified complex is nevertheless completely dependent on Snf7. This apparent paradox can be nicely resolved by our assumption that ESCRT-III formation is initiated by Snf7 and that after completion of the ESCRT-III function, Snf7 is replaced by Mos10 followed by disassembly of the complex.

This notion is supported by fluorescence microcopy of Mos10-sfGFP structures. We could show that the localization of Mos10-sfGFP to endosomal structures at the vacuole is completely dependent on the “upstream” ESCRT-III proteins Vps20, Snf7, Vps2 and Vps24 (Fig. S8-1). In the respective deletion mutants, Mos10-sfGFP was completely cytosolic. By double-labeling experiments, we could further show that the Mos10-sfGFP patches at the vacuole also contain Vps2 and Vps24 (Fig. S8-2). This is in line with the Ni-NTA pulldown experiments. A small discrepancy was noted with respect to Did2. While Did2-mCherry could be clearly detected in the Mos10-sfGFP patches at the vacuole, only a very faint Did2 signal could be obtained in the Ni-NTA pulldown. But this discrepancy could be related to the different behavior of native vs. tagged protein. In the Mos10-sfGFP Snf7-mCherry strain, the Mos10 signal at the vacuole was lost. Instead, Mos10-sfGFP now localized to the bright Snf7-mCherry clusters and to smaller vesicles. Thus, Snf7-mCherry tagging leads to the same phenotype as *VPS68* deletion. This indicates that Snf7 and Vps68 act together at the same step. Also of note, the phenotype of Snf7 tagging was different from the phenotype of the *SNF7* deletion. This shows that Snf7-mCherry is not a completely non-functional protein, but that it preserves part of the Snf7 function.

### Membrane topology of Vps68

Vps68 is a membrane protein with four predicted transmembrane helices (TM). At least one of the four helices (helix 2) appears to be an amphipathic helix with an asymmetric distribution of hydrophilic and hydrophobic amino acids and a high hydrophobic moment (<μ_H_>) (Fig. 9). Vps68 resembles Nce102 in its structure. Nce102 is localized to plasma membrane invaginations called eisosomes. It has an unusual membrane topology with four potential TMs, where only TM1 and TM4 are spanning the membrane, while TM2 and TM3 are localized to the external space (Loibl *et al*, 2010). To see, whether Vps68 has a similar topology, a membrane topology analysis similar to the one described for Nce102 was performed.

**Figure 9.**
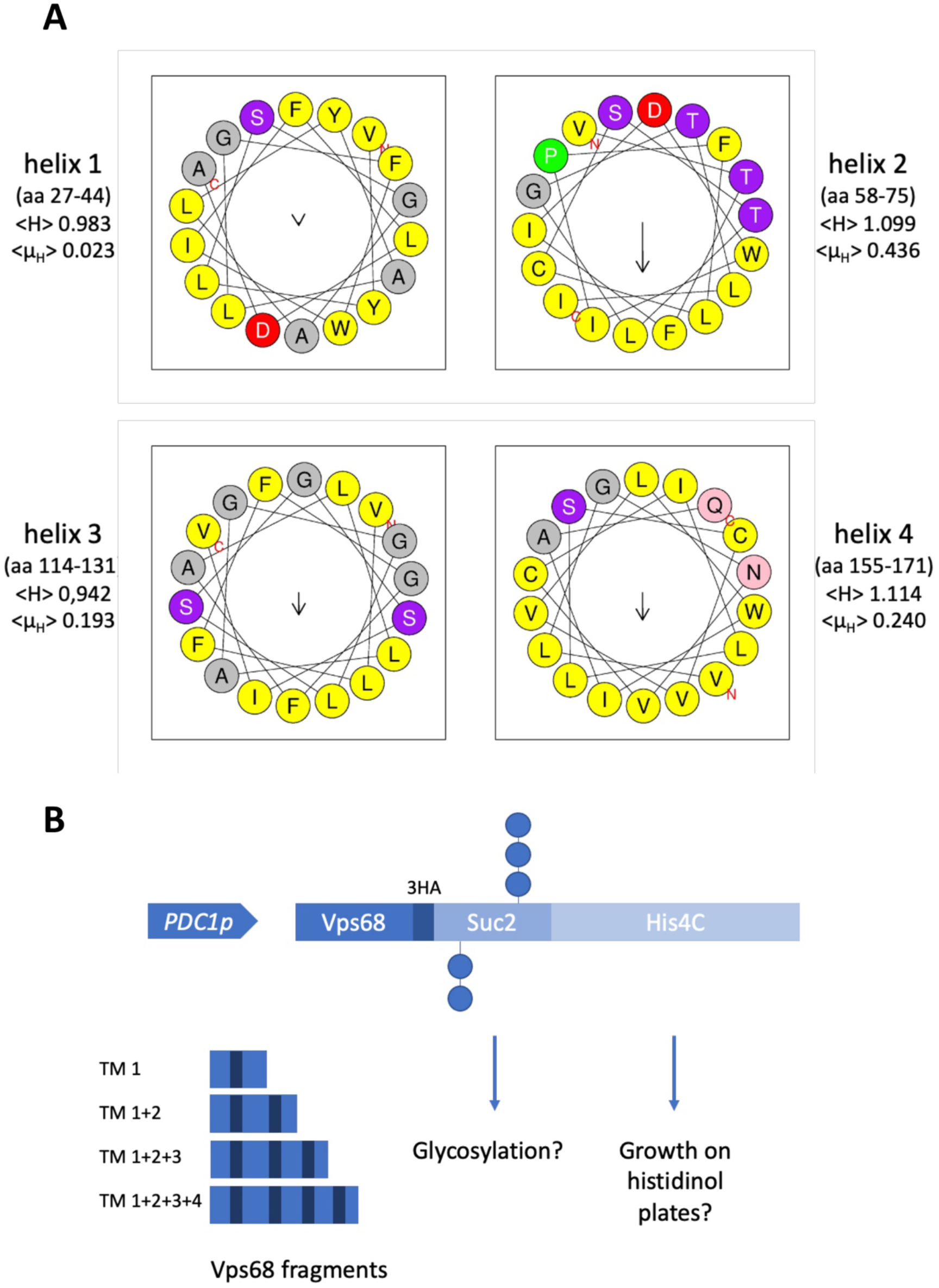
(A) Helical wheel projection of the predicted TMs of Vps68. The predicted TMs of Vps68 were analyzed with HeliQuest (Gautier *et al*, 2008). The hydrophobicity (<H>) and the hydrophobic moment (<μ_H_>) are indicated for each α-helix. (B) Vps68 topology reporter. A schematic representation of the fusion proteins used to study the Vps68 topology (see text).

To probe the topology of Vps68, different N-terminal fragments of Vps68 were fused to the first 307 amino acids of mature invertase (Suc2) followed by His4 deleted for the first 33 amino acids (His4C) (Sengstag *et al*, 1990). Three HA tags were inserted between the Vps68 and the Suc2 portions for western blot detection. The fusion proteins were expressed from single copy plasmids under the control of the *PDC1* promoter (Fig. 9). Depending on the TMs present in the N-terminal Vps68 fragment, the downstream sequences either point to the lumen of the ER or to the cytosol. When they point to the lumen of the ER, N-glycosylation sites in Suc2 are modified by the attachment of sugar chains, which leads to a mobility shift of the fusion protein on SDS gels. To prove that the mobility shift is caused by glycosylation, the sugar chains can be removed by Endoglycosidase H (Endo H) treatment. The truncated His4C protein has histidinol dehydrogenase activity. Yeast cells expressing this protein are able to grow on histidinol plates in the absence of histidine in a *HOL1-1* strain background (Sengstag *et al*., 1990). But growth is only possible, when His4C is present in the cytosol. Thus, we have two complementary ways of assessing the localization of the sequences downstream of the Vps68 fragments.

The western blot analysis of the different fusion constructs is presented in Fig. 10A. The fusion protein without Vps68 sequences runs at the expected position of 114,3 kDa and is not affected by Endo H treatment. Insertion of the N-terminal Vps68 fragment containing TM1 leads to a shift to higher mobility, larger than the one expected from the addition of the Vps68 sequences. This shift is reversed by Endo H treatment. This shows that TM1 spans the membrane and that the N-terminus of Vps68 points to the cytosol, while the rest of the fusion protein is inside the ER. When a fragment containing TM1 and TM2 is inserted, the same pattern is obtained. This shows that TM2 does not span the membrane and that the sequences downstream of TM1 are still oriented towards the lumen of the ER. This fusion protein appears to be unstable, because the western blot signal is weaker than the signal for the previous constructs. The same pattern was observed with the Vps68 fragment containing TM1, TM2 and TM3, i.e. TM3 also does not span the membrane. Finally, when the complete sequence of Vps68, containing all four TMs, was inserted, no Endo H dependent mobility shift was observed. This indicates that TM4 traverses the membrane and that the C-terminus of Vps68 points to the cytosol. Now, the same strains were streaked out on plates containing either histidine or histidinol (Fig. 10B). While all strains grew on the histidine plate, only the fusion with full-length Vps68 and the control without Vps68 sequences were able to grow on the histidinol plate, i.e. in these constructs the His4C part is localized to the cytosol. This result is the exact mirror image of the western blot experiment. Taken together, this experiment demonstrates that Vps68 has indeed the same topology as Nce102. Vps68 forms a hairpin structure with TM1 and TM4 traversing the membrane and TM2 and TM3 being localized to the luminal side of the endosomal membrane (Fig. 10C). Both TM2 and TM3 appear to be amphipathic helices that are able to interact with the luminal leaflet of the endosomal membrane.

**Figure 10.**
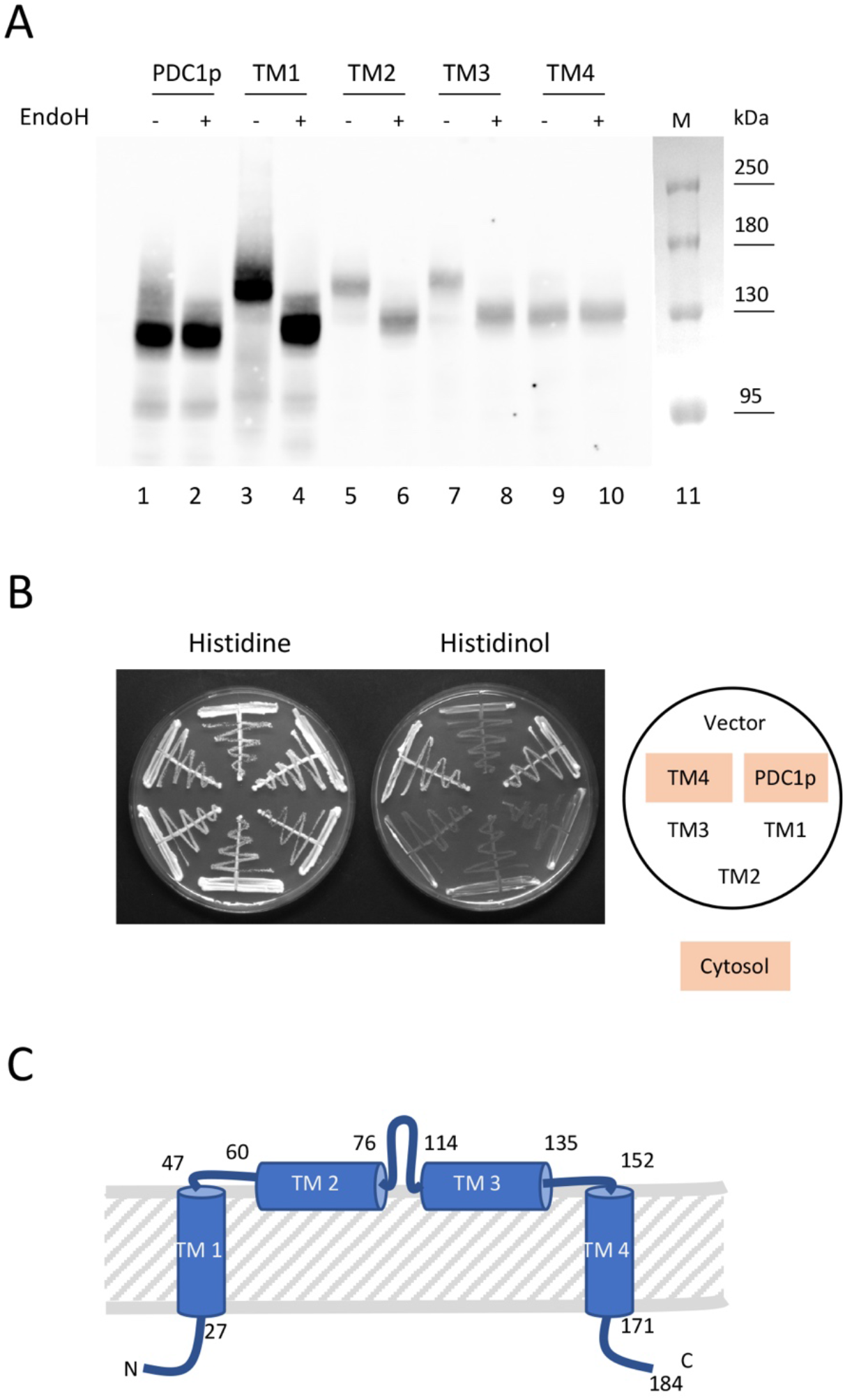
Membrane topology of Vps68. The *HOL1-1* strain STY50 was transformed with plasmids expressing different fragments of Vps68 fused to Suc2 and His4C fragments. (A) Glycosylation patterns of fusion proteins. Cell extracts of the transformants were examined for the fusion proteins by western blotting with anti-HA antibodies. Cell extracts were either treated with Endo H to remove N-glycosylation (lanes with even numbers, Endo H +) or were left untreated (lanes with odd numbers, Endo H -). The following plasmids were used: pRK2015 (lanes 1,2, no Vps68), pRK2016 (lanes 3,4, Vps68 with TM1), pRK2017 (lanes 5,6, Vps68 with TM1+TM2), pRK2018 (lanes 7,8, Vps68 with TM1+TM2+TM3) and pRK2019 (lanes 9,10, full-length Vps68). Lane 11: protein marker. (B) Growth on histidinol plates. The transformants were plated on agar plates containing histidine (left) or histidinol (right). The plasmids contained in the transformants were (starting from the top, counterclockwise direction): YCplac33 (vector), pRK2015, pRK2016, pRK2017, pRK2018, pRK2019. The plasmids expressing fusion proteins, whose C-terminus points to the cytosol are highlighted. (C) Membrane topology predicted from the experiments. The positions of the TMs in the Vps68 sequence are indicated.

## Discussion

Here we present evidence that Vps68 interacts with ESCRT-III and that it cooperates with ESCRT-III in ILV formation and abscission. Our data further suggest that at some point in the ESCRT-III functional cycle Snf7 is replaced by Mos10. This switch in ESCRT-III subunits could be associated with a transition from an active complex to a disassembly competent complex.

### Vps68 interacts with ESCRT-III

In a previous report, it was proposed that Vps68 acts with or downstream of the ESCRT machinery to regulate a novel step in endosome biogenesis (Schluter *et al*., 2008). Our findings now suggest that Vps68 is tightly connected to ESCRT-III function. First of all, we were able to show that Vps68 physically interacts with ESCRT-III. At present it is not clear how Vps68 interacts with ESCRT-III, but Mos10 seems to be dispensable for the interaction, since ESCRT-III subunits could still be immunoprecipitated by Vps68 in the absence of Mos10. Loss of Vps68 leads to a shift in the composition of ESCRT-III. The number of core components in the complex (Snf7, Vps2 and Vps24) increases, while the number of the associated subunits (Did2, Ist1 and Mos10) decreases. This supports the view that at some point during the functional cycle a switch occurs from an active complex involved in ILV formation and/or abscission to a disassembly competent complex. This switch is reflected in a change in ESCRT-III composition. The precise function of ESCRT-III during the formation of an intraluminal vesicle is still not clear. ESCRT-III could be involved in the deformation of the endosomal membrane, or its function could be restricted to the release of the vesicle from the membrane into the lumen of the endosome. In any case, these events have to be tightly coordinated with the incorporation of cargo proteins into the forming vesicle. The function of ESCRT-III is completed, when all cargo proteins are removed from the endosomal membrane. The cue that signals completion of the ESCRT-III task is not known, but it is tempting to speculate that the presence of ubiquitinated cargo proteins in the endosomal membrane is part of the signal. As soon as the ubiquitinated cargos are removed, ESCRT-III would switch to a disassembly competent complex followed by its rapid dissolution. The ESCRT-III associated subunits have been implicated in ESCRT-III disassembly (Azmi *et al*., 2008; Dimaano *et al*., 2008; Rue *et al*., 2008). This notion is reinforced by our observation that ESCRT-III core components accumulate at the membrane in a *MOS10* deletion. The finding that the associated subunits are underrepresented in the *Δvps68* strain, while the core subunits are overrepresented, indicates that the completion of the ESCRT-III task is delayed in *Δvps68*. A close connection between the Vp68 function and the presence of cargo protein is also suggested by our epistasis analysis. When the cargo protein Ste6-sfGFP was expressed to near wildtype levels, an effect of the *VPS68* deletion on the localization Ste6-sfGFP was barely detectable. But upon overexpression, a class E-like phenotype was observed. From this we conclude that Vps68 assists ESCRT-III in its function. When the load on the endocytic pathway is high, Vps68 is especially critical for the function of the ESCRT-III system.

### Role of Vps68

What could be the role of Vps68? A potential function is suggested by the unusual membrane topology of Vps68. We found that the two potential membrane helices 2 and 3 are not spanning the membrane, but are instead localized to the luminal side of the endosomal membrane. At least helix 2 appears to be an amphipathic helix (AH) with a hydrophilic and a hydrophobic face and a high hydrophobic moment. Helix 3 is probably also an amphipathic helix. Amphipathic helices have the tendency to interact with membrane surfaces. A number of functions have been ascribed to amphipathic helices (Gimenez-Andres *et al*, 2018). The AH containing protein Pex11, for instance, deforms membranes and induces tubulation. Some AHs can sense or stabilize membrane curvature, they act as amphipathic lipid packing sensors (ALPS). Yet other AHs like in melittin disturb the integrity of the lipid bilayer. We propose that Vps68, analogous to melittin, facilitates ILV abscission by disturbing the order of the lipids in the luminal leaflet of the bilayer. ESCRT-III in turn could act in the same way on the cytosolic leaflet. By simultaneously acting on opposing leaflets of the bilayer, ESCRT-III and Vps68 could promote the release of the ILV from the endosomal membrane.

### Switch from a Snf7 to a Mos10 containing ESCRT-III complex

As described above, it appears plausible that during an ESCRT-III functional cycle a shift occurs from an active complex to a disassembly competent complex. The shift is accompanied by a change in ESCRT-III composition. We think, the key event in this transition is the replacement of Snf7 by Mos10. This conclusion is derived from our co-immunoprecipitation and pulldown experiments. In Ni-NTA pulldowns with Mos10-6His, the core ESCRT-III subunits Vps2 and Vps24 were co-purified (and a small amount of Did2), but no Snf7. In principle, this Mos10 complex could be unrelated to the endosomal ESCRT-III complex. But we think this is not the case. We found that the co-purification of ESCRT-III subunits with Mos10 was completely Snf7 dependent. Our interpretation of this finding is that Snf7 initiates ESCRT-III formation, but that it is replaced later on by Mos10. Snf7 and Mos10 are closely related. The ESCRT-III proteins can be divided into two classes, based on sequence homology (Leung *et al*., 2008). A member of each class was already present in the last common eukaryotic ancestor (LCEA). The first class consists of Snf7, Mos10 and Vps20 (and also Chm7) and the second class contains Did2, Vps2 and Vps24. In fact, in our initial report, we were only able to identify the first class of ESCRT-III proteins, the second class escaped our attention due to divergence in the primary sequence (Kranz *et al*., 2001).

A number of observations suggest that Mos10 acts after Snf7. A careful study of the morphology of MVBs in ESCRT-III mutants showed a clear distinction between the core components Snf7, Vps2, Vps20 and Vps24 and the ESCRT-III associated proteins Did2 and Mos10 (Nickerson *et al*., 2010). When one of the core components was deleted, no ILVs were formed. Instead, the typical class E structures, closely juxtaposed flatted endosomes could be observed. In *did2* and *mos10* mutants, ILVs were stilled formed, but the MVBs assumed an elongated morphology called “vesicular tubular endosomes” VTEs. Previous evidence and the findings in our study clearly point to a role of Mos10 in the disassembly of ESCRT-III, which appears to occur after ILV formation is finished.

### Licensing in the endocytic pathway

Mos10-sfGFP shows a unique intracellular distribution. While the other ESCRT-III proteins localized to punctate structures, Mos10-sfGFP was detected at the vacuolar membrane. For the localization studies, we used C-terminally tagged ESCRT-III proteins. Most of these tagged proteins were compromised in their function. But still, useful information can be gained from the investigation of these proteins. The phenotypes of the tagged proteins were clearly different from complete knockouts, which shows that the function of these proteins is partly preserved. In fact, we think that these proteins can be used as tools to dissect the ESCRT-III cycle, since tagging of different ESCRT-III proteins arrests the cycle at distinct steps. Thus, tagging of Snf7 leads to an early arrest, while tagging of Mos10 arrests the cycle at a later time point. It appears that an early arrest, like the one imposed by tagging of Snf7, prevents downstream events like tethering and fusion of late endosomes with the vacuolar membrane. For this reason, the tagged core ESCRT-III proteins were detected in endosomal structures and not at the vacuole. The successive events in the endocytic pathway seem to be strictly depended on each other. Only upon successful completion of one step, the next step is licensed. Thus, progression to endosome tethering only occurs when ILV formation is completed. Disassembly of ESCRT-III does not seem to be a precondition for endosome tethering to the vacuole. That is why the ESCRT-III complex containing Mos10-sfGFP is detected in endosomal structures tethered to the vacuole. ESCRT-III disassembly in turn could be a precondition for endosome fusion, since we observed a large number of Mos10-sfGFP structures at the vacuolar membrane. Mos10-sfGFP was not detected at the vacuolar membrane in a strain expressing tagged Snf7 or in a strain carrying a *VPS68* deletion. In these strains, Mos10-sfGFP showed an endosomal localization. This further supports the idea that Snf7 and Vps68 closely act together and that progression to endosome tethering is blocked, when these functions are compromised.

## Materials and methods

### Plasmids and yeast strains

Yeast cells were grown in YPD medium (1 % yeast extract, 2 % peptone, 2 % glucose) for immunoprecipitation experiments and in SD/CAS medium (0.67 % yeast nitrogen base, 1 % casamino acids, 2 % glucose, 50 mg/l uracil and tryptophan) for fluorescence microscopy. For SILAC, yeasts were grown in SD medium (0.67 % yeast nitrogen base, 2 % glucose) with the required auxotrophic markers. Arginine and lysine labeled with heavy isotopes [13C6/15N2] were obtained from Silantes GmbH (Munich, Germany). The yeast strains used are listed in Tab. S2. All yeast strains constructed in this study are derived from JD52 (J. Dohmen, Cologne, Germany) by the integration of PCR-cassettes into the yeast genome (Longtine *et al*, 1998). Some deletions were generated by the CRISPR/Cas9 technique (Brune *et al*, 2019). The plasmids used in this study are listed in Tab. S3.

### Mos10-6His purification by immobilized metal affinity chromatography (IMAC)

For SILAC (stable isotope labeling with amino acids in cell culture) experiments, cells were grown in SD medium with 50 mg/l adenine, histidine, uracil, tryptophan, 120 mg/l leucine and 60 mg/l arginine and lysine. Cells were grown overnight to exponential phase (OD_600_ < 1.0) at 30°C, harvested and washed with PBS. The wildtype strain RKY2998 was grown in light medium and the Mos10-6His strain RKY2999 was grown in heavy medium. Equal amounts of the heavy and light cells were mixed and resuspended in 10 ml PBS. The cells were lysed by glass-beading in a Fast Prep24 machine (MP Biomedicals, Eschwege, Germany) (2-times, 1 min, 4 m/s). Next, the cell lysate was incubated with 4 mM disuccinimidyl suberate (DSS) crosslinker for 1 h at 4°C on a rocker. The reaction was stopped by quenching with 100 mM

Tris-HCl pH 8 and the membranes were solubilized with 1 % Triton X-100 for 15 min at 4°C. To remove cell debris, the lysate was spun at 15,000 g at 4°C for 15 min. The supernatant was adjusted to 300 mM NaCl and 15 mM imidazole and filtered through a 0.45 μm filter. The sample was then applied to a AEKTA start chromatography system (GE Healthcare Europe GmbH, Freiburg, Germany), and purification of crosslinked proteins was achieved on a 1 ml HisTrap™ Fast Flow column. Bound proteins were eluted from the column by a 50-500 mM imidazole gradient. The collected fractions were analyzed by SDS-PAGE with Coomassie blue staining and western blotting with anti-Mos10 antibodies. The fractions containing Mos10-6His crosslinks were analyzed by mass spectrometry (Core Facility, University of Hohenheim). Mos10-6His was also purified by IMAC on a small scale. Cells were grown in YPD medium to exponential phase and 20 OD_600_ of cells were harvested. Basically, the cells were processed as described below for the co-immunoprecipitation with the modification that after the 500 g spin, 20 mM imidazole and 50 μl of a 50 % Ni-NTA bead slurry were added instead of the antibodies and protein A sepharose beads.

### Co-immunoprecipitation

For Co-immunoprecipitation (Co-IP) experiments, 50 OD_600_ cells of an exponential YPD culture were harvested, washed with PBS and resuspended in 200 μl PBS with protease inhibitors. The cells were lysed by glass-beading for 5 min at 4°C. After lysis, 600 μl PBS and 2.5 mM of the cleavable crosslinker dithiobis (succinimidylpropionate) (DSP) were added. The solution was incubated for 1 h at 4 °C on a rocker. The crosslinking reaction was stopped by quenching with 100 mM Tris-HCl pH 8.0 and the membranes were dissolved by 1 % Triton X-100. The reaction was incubated for 15 min at 4 °C on a rocker and then centrifuged for 10 min at 500 g at 4°C to remove cell debris. Then the supernatant was incubated with 10 μl antiserum for 1 h at 4°C on a rocker followed by an incubation with 80 μl of a 50 % slurry of protein A sepharose beads for 1 h at 4°C. The solution was washed three times with 1 ml PBS at 100 g for 30 s. The beads were resuspended in a mixture of 100 μl PBS and 100 μl SDS sample buffer and heated to 95°C for 5 min.

### Endoglycosidase H treatment

The cells were grown to exponential phase (OD_600_ < 1) and 10 OD_600_ were harvested, washed in 10 mM NaN_3_ and resuspended in 100 μl of lysis buffer (0.3 M sorbitol, 50 mM HEPES pH 7.5) with protease inhibitors. After glass-beading for 5 min at 4°C, 150 μl of SDS sample buffer were added and then the mixture was heated at 50°C for 20 min. The lysate was centrifuged at 20,000 g for 5 min in a table top centrifuge to remove cell debris. The supernatant was diluted 1:10 with Endo H buffer (50 mM Tris-HCl pH 6.8, 50 mM EDTA, 1 % Triton X-100). Two 100 μl aliquots were incubated overnight at 37°C with and without Endo H (3000 U, NEB). Then the samples were heated with 100 μl SDS sample buffer for 5 min at 95°C.

### Cell fractionation

The sucrose gradients were performed as described in (Heinzle *et al*., 2019). Flotation was performed as described in (Pawelec *et al*, 2010).

## Acknowledgements

We thank Roger Schneiter for sending us the *HOL1-1* strain STY50. This project was funded by the German Research Fund (DFG) KO 963/8-1.

## Supplemental figures

**Figure S2.**
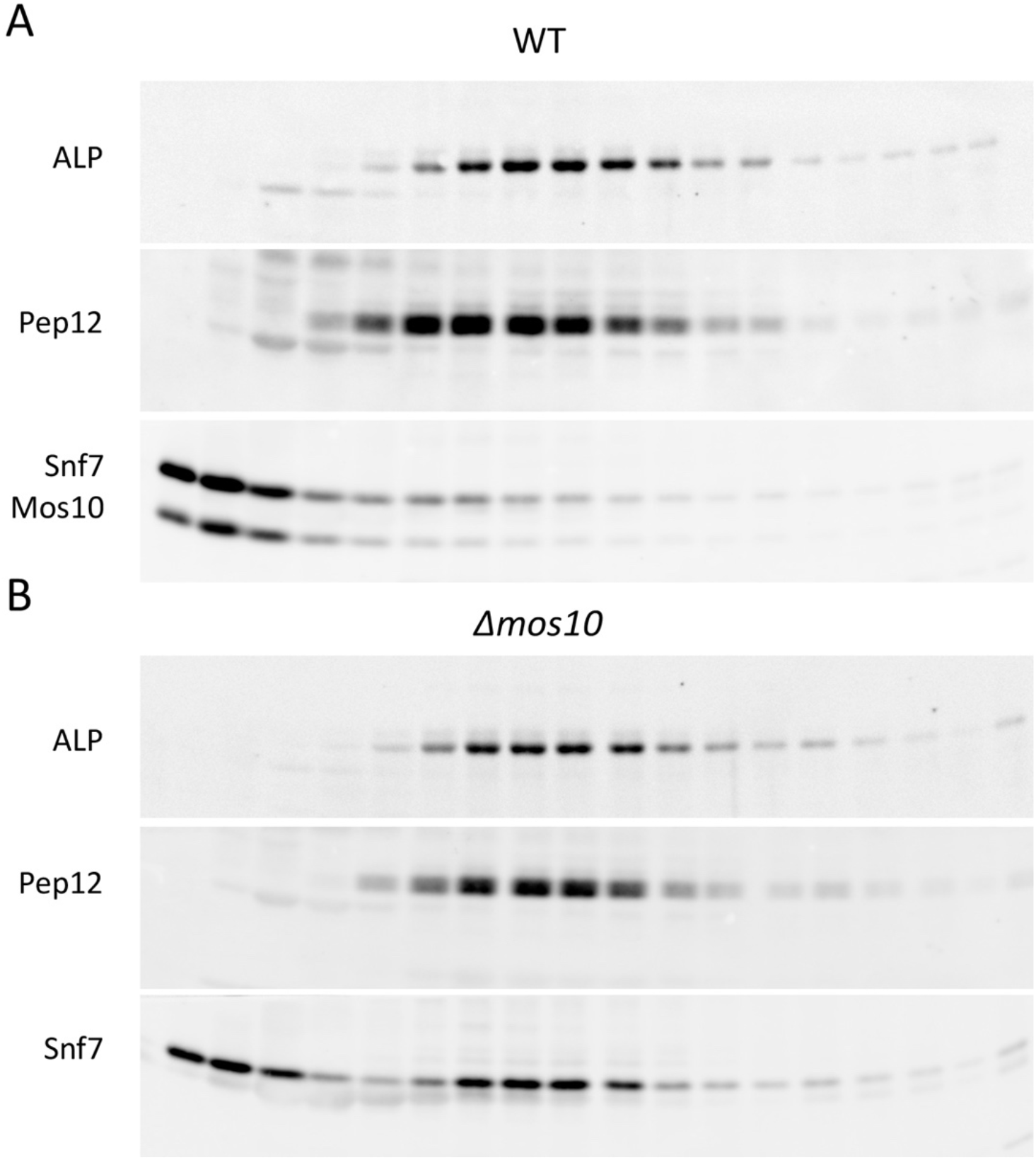
Membrane accumulation of Snf7 in a *MOS10* deletion mutant. Western blots to Fig. 2. (A) RKY1558 (WT), (B) RKY2909 (*Δmos10*), proteins (from top to bottom): alkaline phosphatase (ALP), Pep12, Snf7/Mos10.

**Figure S3.**
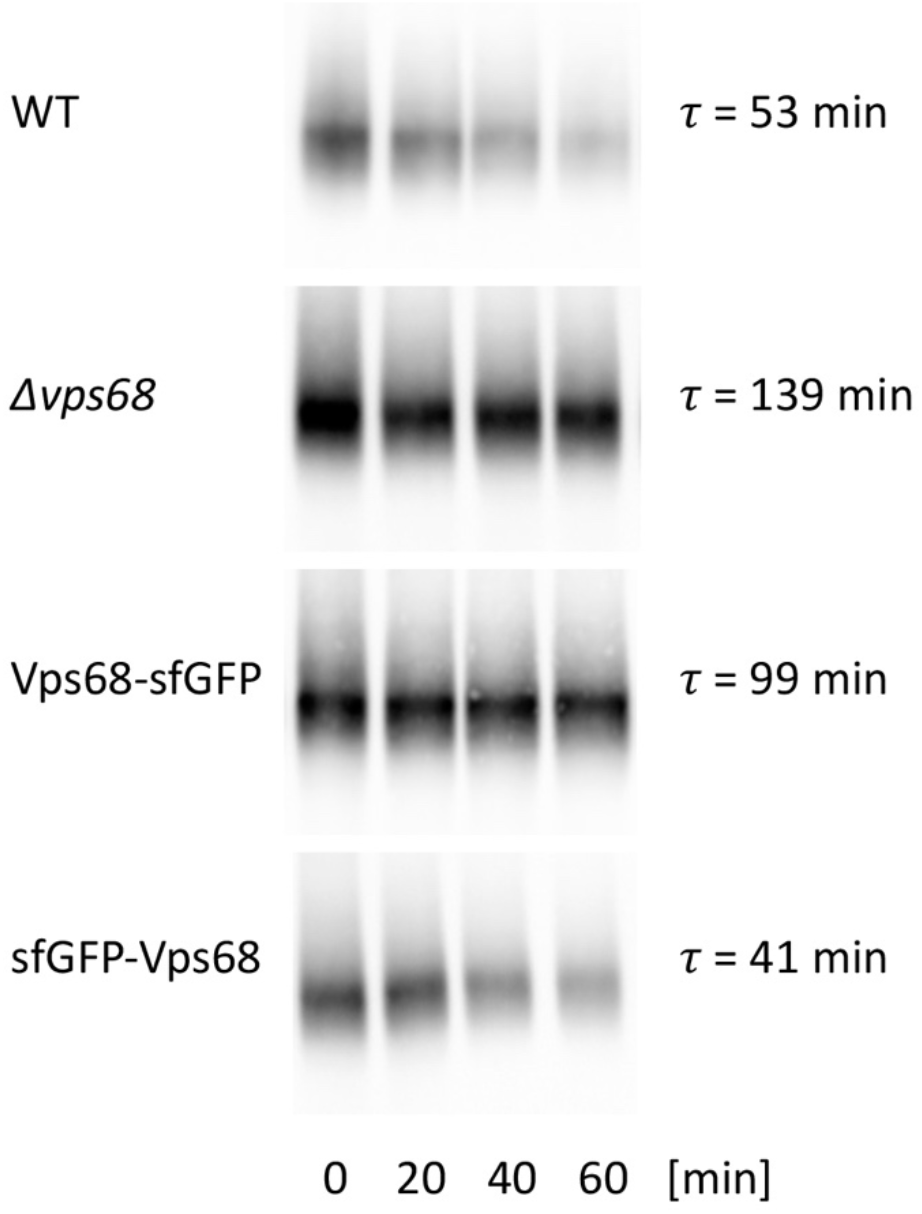
Ste6 turnover. Cycloheximide chase. Yeast cells were grown in YPD medium. At t_0_ 100 μg/ml cycloheximide were added. Cell extracts of equal volumes of the culture were prepared at the times indicated and examined for Ste6 by western blotting. Strains (from top to bottom): RKY1558 (wildtype), RKY3222 (*Δvps68*), RKY3183 (*VPS68-sfGFP*), RKY3285 (*sfGFP-VPS68*). The signals were quantified by ImageJ and the Ste6 half-lives were calculated (as indicated). Ste6 half-lives were higher with CHX than in the gal depletion experiment, due to inhibitory effects of CHX on endocytic trafficking.

**Figure S4.**
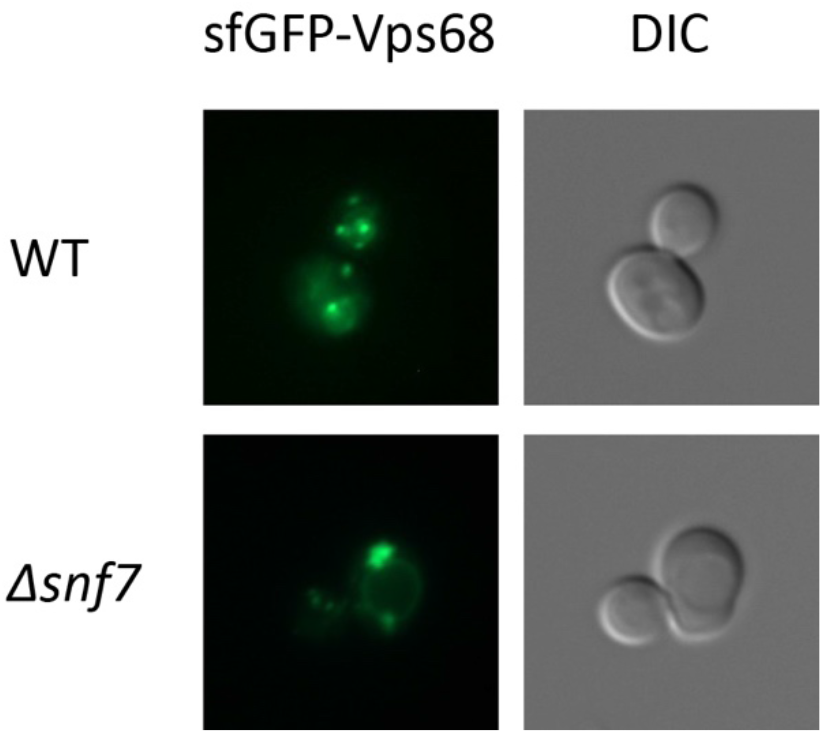
Localization of sfGFP-Vps68. Strains, top: RKY3285 (*sfGFP-VPS68*), bottom: RKY3412 (*sfGFP-VPS68 Δsnf7*). Left panels: sfGFP-Vps68 fluorescence, right panels: DIC image.

**Figure S5.**
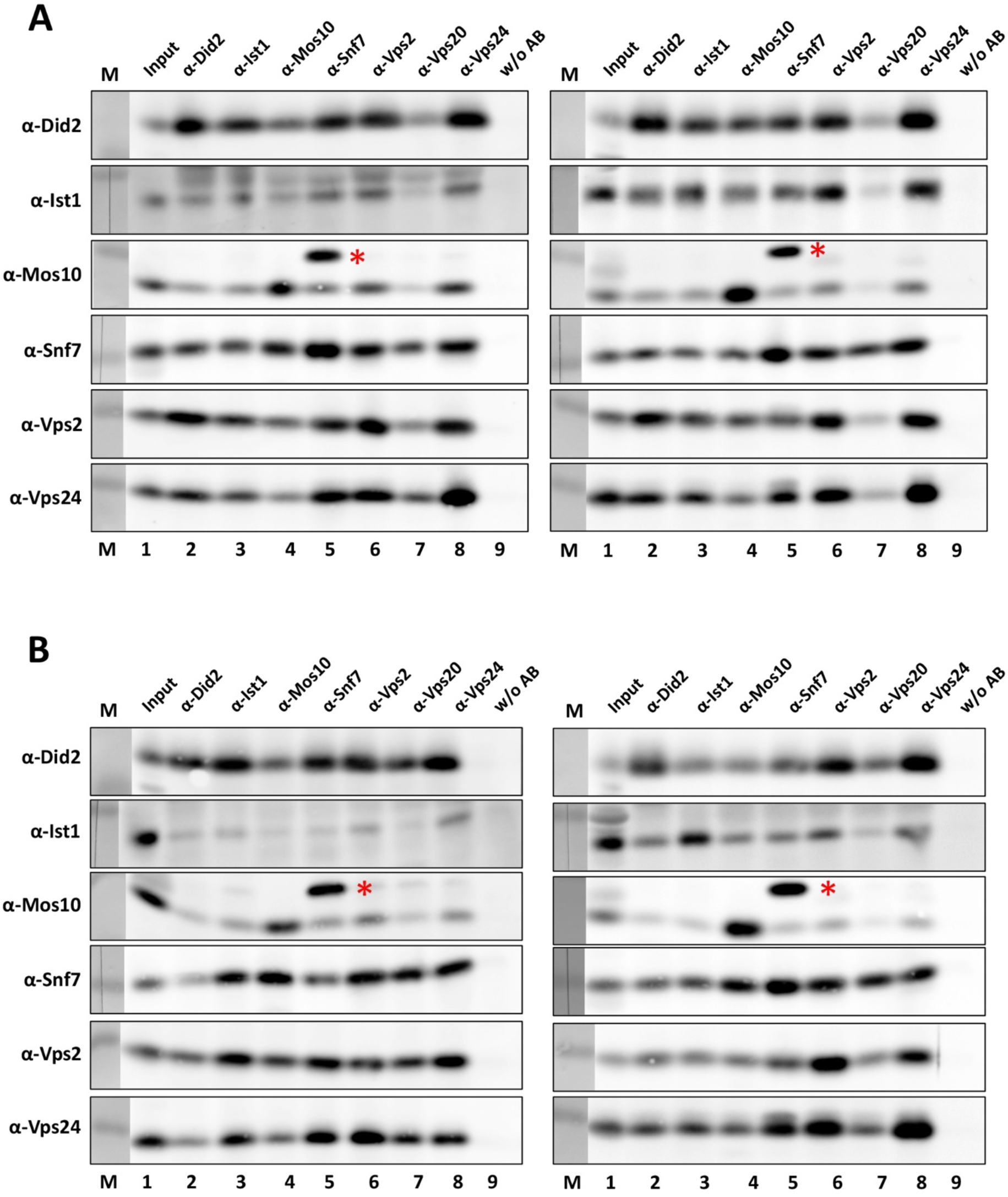
Co-immunoprecipitation of ESCRT-III proteins. ESCRT-III proteins were immunoprecipitated from cell extracts with specific antibodies. The antibodies used for the primary IP are indicated on top of the diagrams (lanes 2-8). Lane 1: input, lane 9: negative control, IP reaction without antibodies, M: protein marker. The immunoprecipitates were examined for co-immunoprecipitation of other ESCRT-III proteins by western blotting with the antibodies indicated on the left side of the diagrams. (A) JD52 (wildtype), (B) RKY3222 (*Δvps68*), two independent experiments each. *Cross-reactivity of anti-Mos10 with Snf7.

**Figure S6-1.**
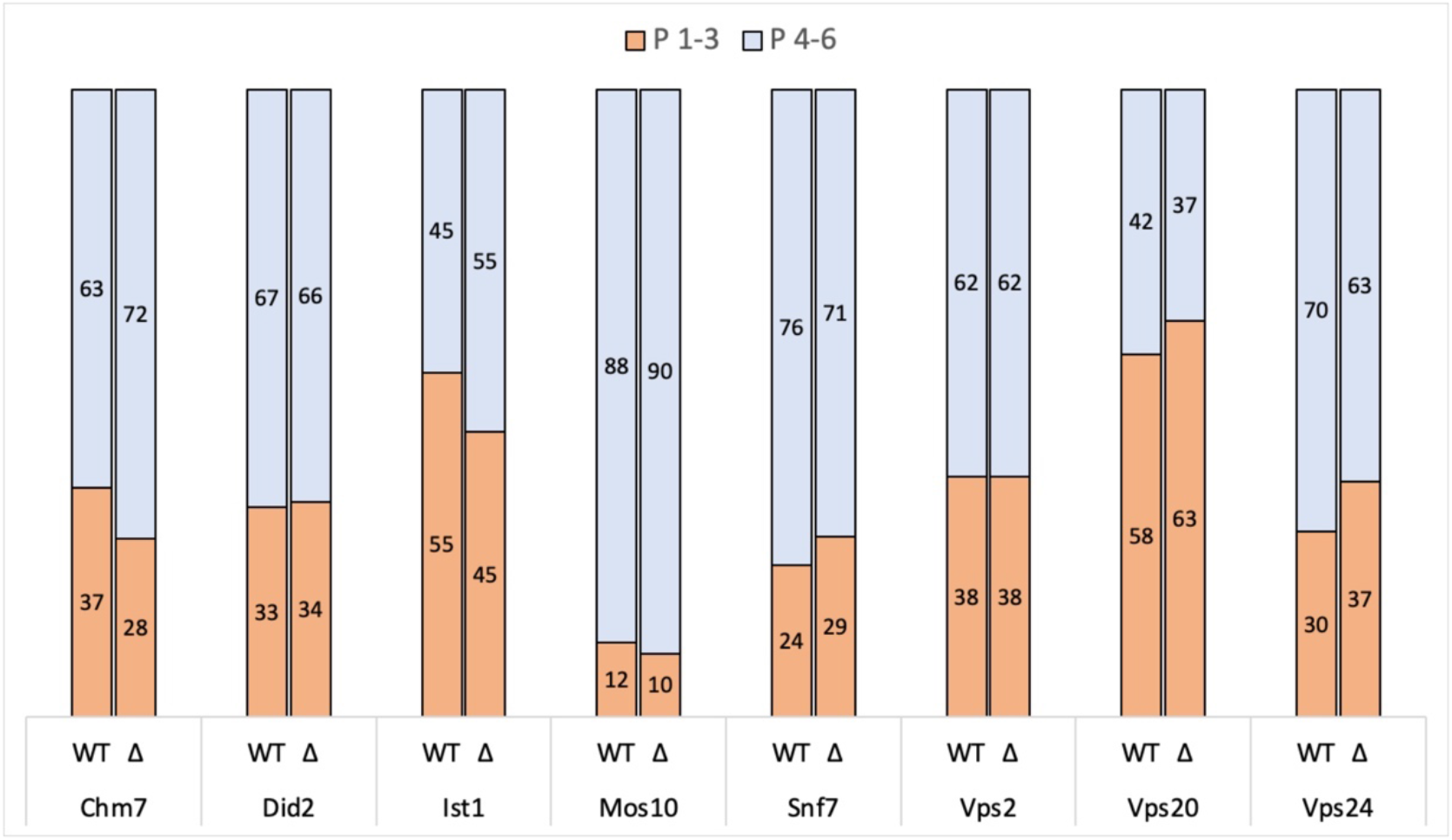
Membrane association of ESCRT-III proteins. The fractions of a flotation gradient were examined for the presence of ESCRT-III proteins by western blotting. The top three fractions of the flotation gradient (P 1-3, orange) contain the membranes and the lower three fractions (P 4-6, blue) contain the soluble proteins. The percentage of the proteins present in these two fractions is indicated. Two columns are shown for each ESCRT-III protein. Left column: extract from the wildtype JD52 (WT), right column: extract from the *Δvps68* strain RKY3222 (Δ). The ESCRT-III proteins fractionated are indicated below the columns.

**Figure S6-2.**
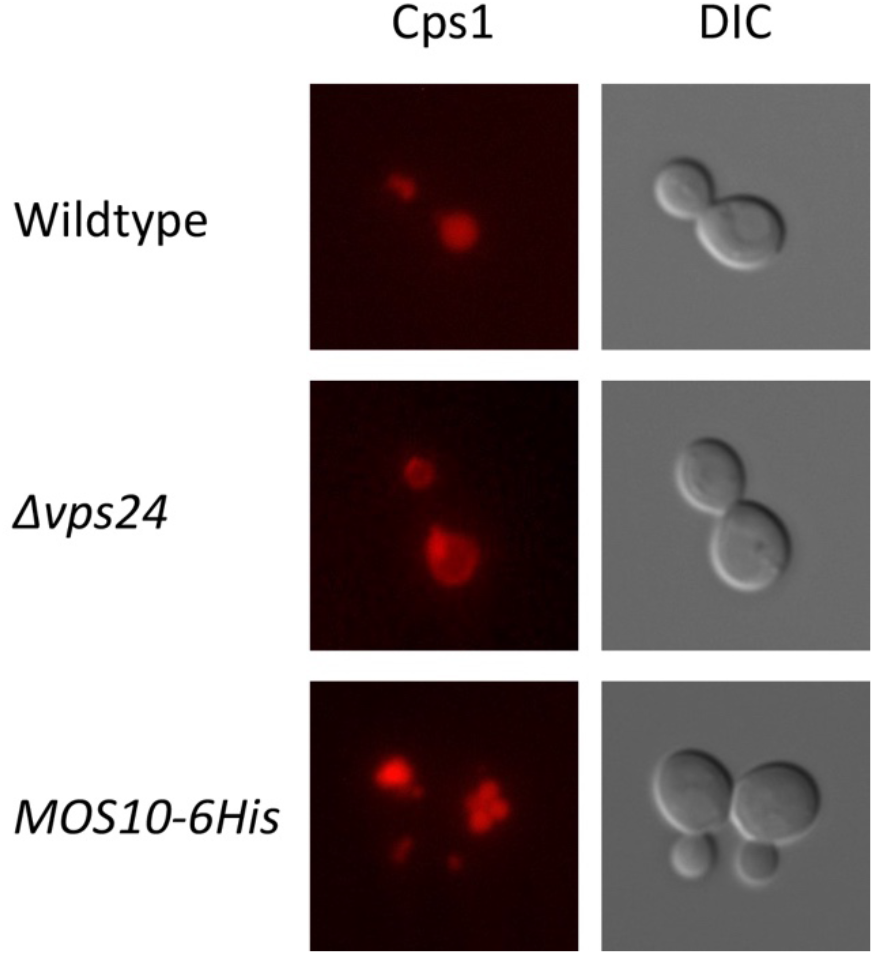
Test for functionality of ESCRT-III-sfGFP fusions, controls. Different yeast strains were transformed with the centromere plasmid pRK1408 expressing mCherry-Cps1 from the *HXT7* promoter. Left panels: mCherry-Cps1 fluorescence, right panels: DIC image. Strains from top to bottom: JD52 (wildtype), RKY2830 (*Δvps24*), RKY2889 (*MOS10-6His*).

**Figure S6-3.**
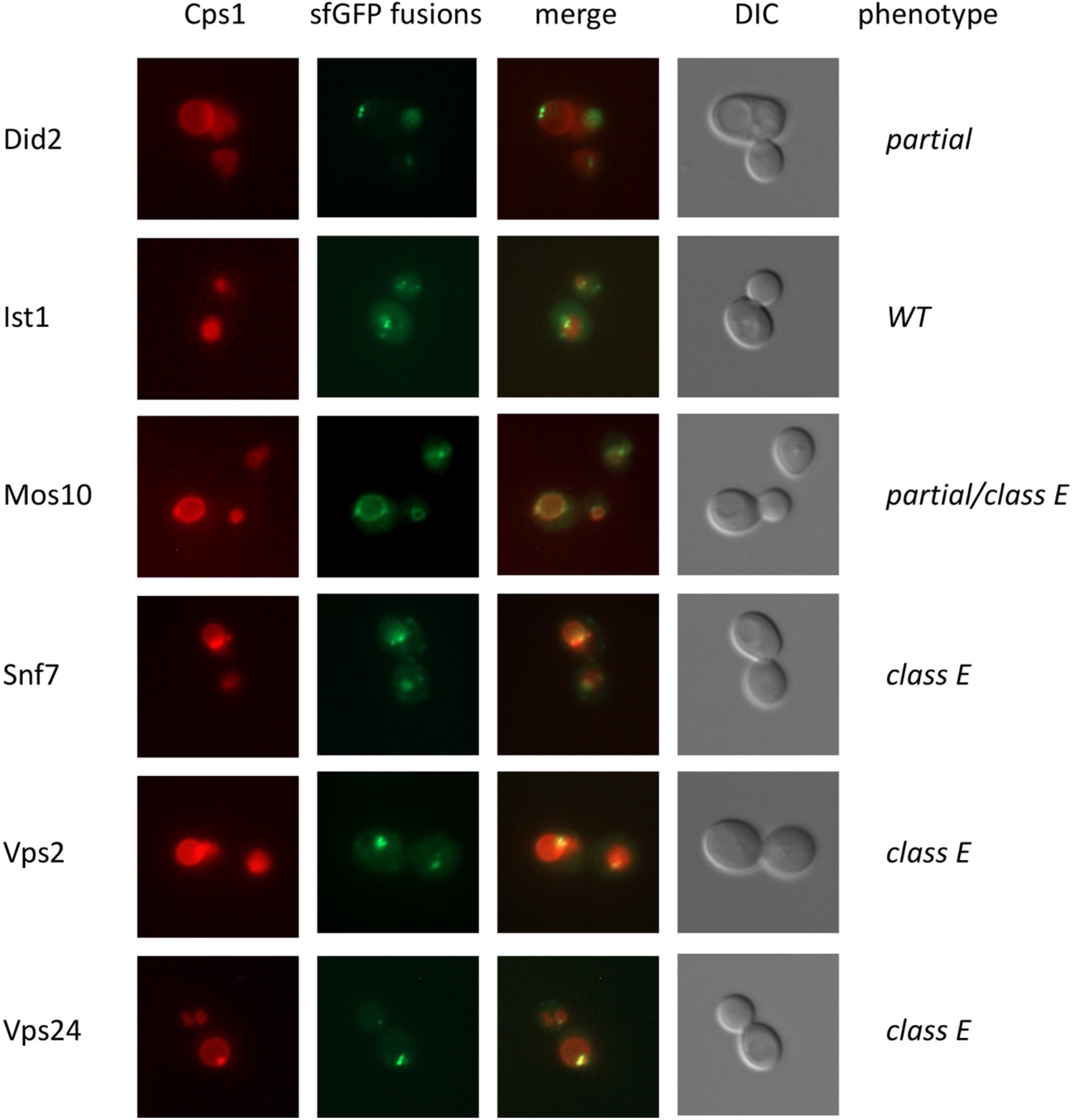
Test for functionality of ESCRT-III-sfGFP fusions. Yeast strains expressing ESCRT-III-sfGFP fusions from their chromosomal loci were transformed with the centromere plasmid pRK1408 expressing mCherry-Cps1 from the *HXT7* promoter. Panels from left to right: mCherry-Cps1 fluorescence, ESCRT-III-sfGFP fluorescence, merged image, DIC image. Strains from top to bottom: RKY3214 (Did2-sfGFP), RKY3215 (Ist1-sfGFP), RKY3216 (Mos10-sfGFP), RKY3217 (Snf7-sfGFP), RKY3218 (Vps2-sfGFP), RKY3220 (Vps24-sfGFP). The Cps1 sorting phenotype is indicated on the right side of the figure.

**Figure S6-4.**
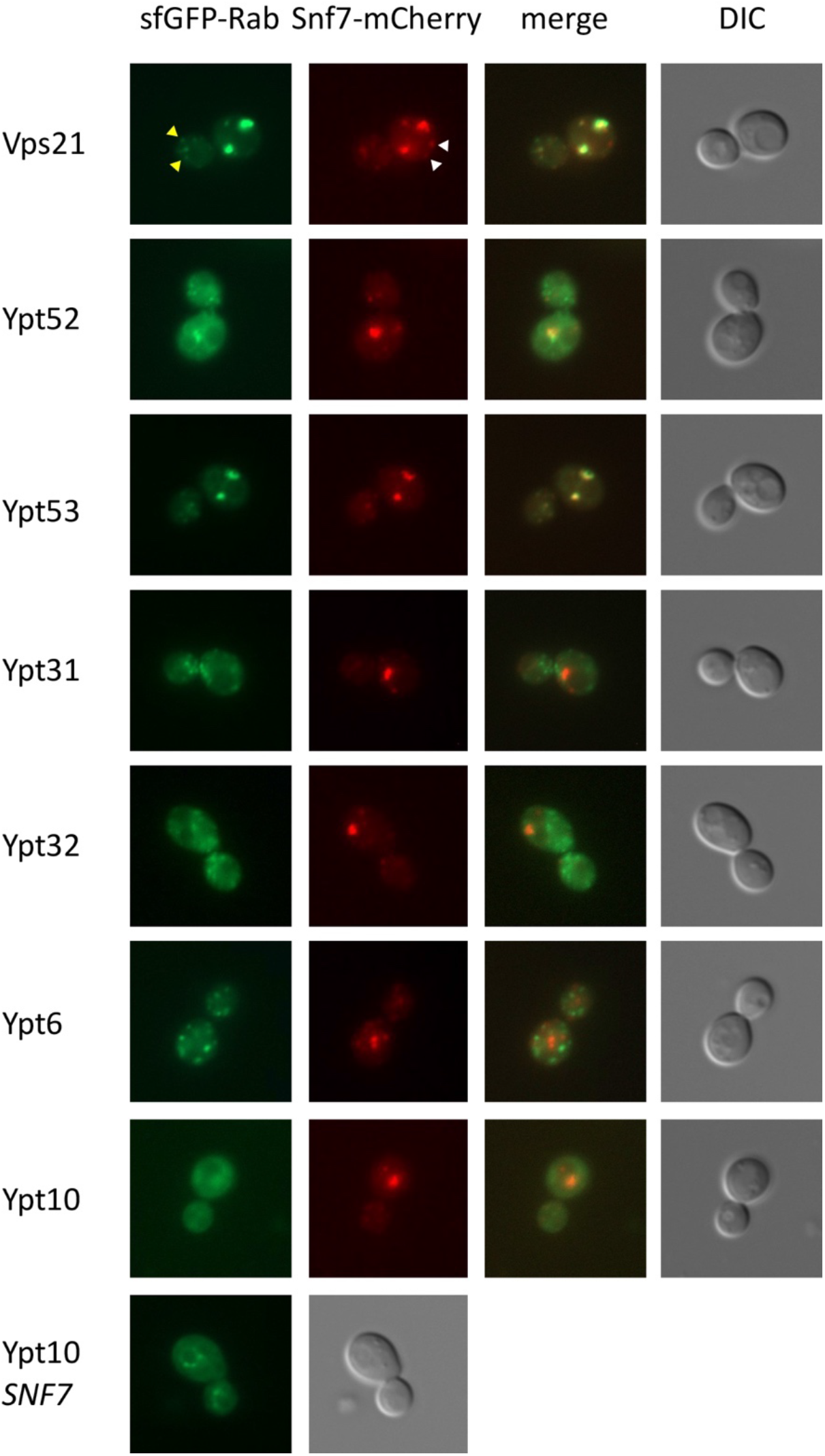
Co-localization of sfGFP-Rab proteins with Snf7-mCherry. Yeast strains expressing N-terminally tagged sfGFP-Rab proteins and C-terminally tagged Snf7-mCherry from their chromosomal loci were examined by fluorescence microscopy. Panels from left to right: sfGFP-Rab fluorescence, Snf7-mCherry fluorescence, merged image, DIC image. Strains from top to bottom (all expressing Snf7-mCherry): RKY3423 (sfGFP-Vps21), RKY3425 (sfGFP-Ypt52), RKY3361 (sfGFP-Ypt53), RKY3362 (sfGFP-Ypt31), RKY3363 (sfGFP-Ypt32), RKY3364 (sfGFP-Ypt6), RKY3365 (sfGFP-Ypt10). Bottom panels: RKY3351 (sfGFP-Ypt10 without Snf7-mCherry, only GFP fluorescence and DIC image are shown). Arrows in Vps21 panel, yellow: sfGFP-Vps21 only, white: Snf7-mCherry only.

**Figure S7.**
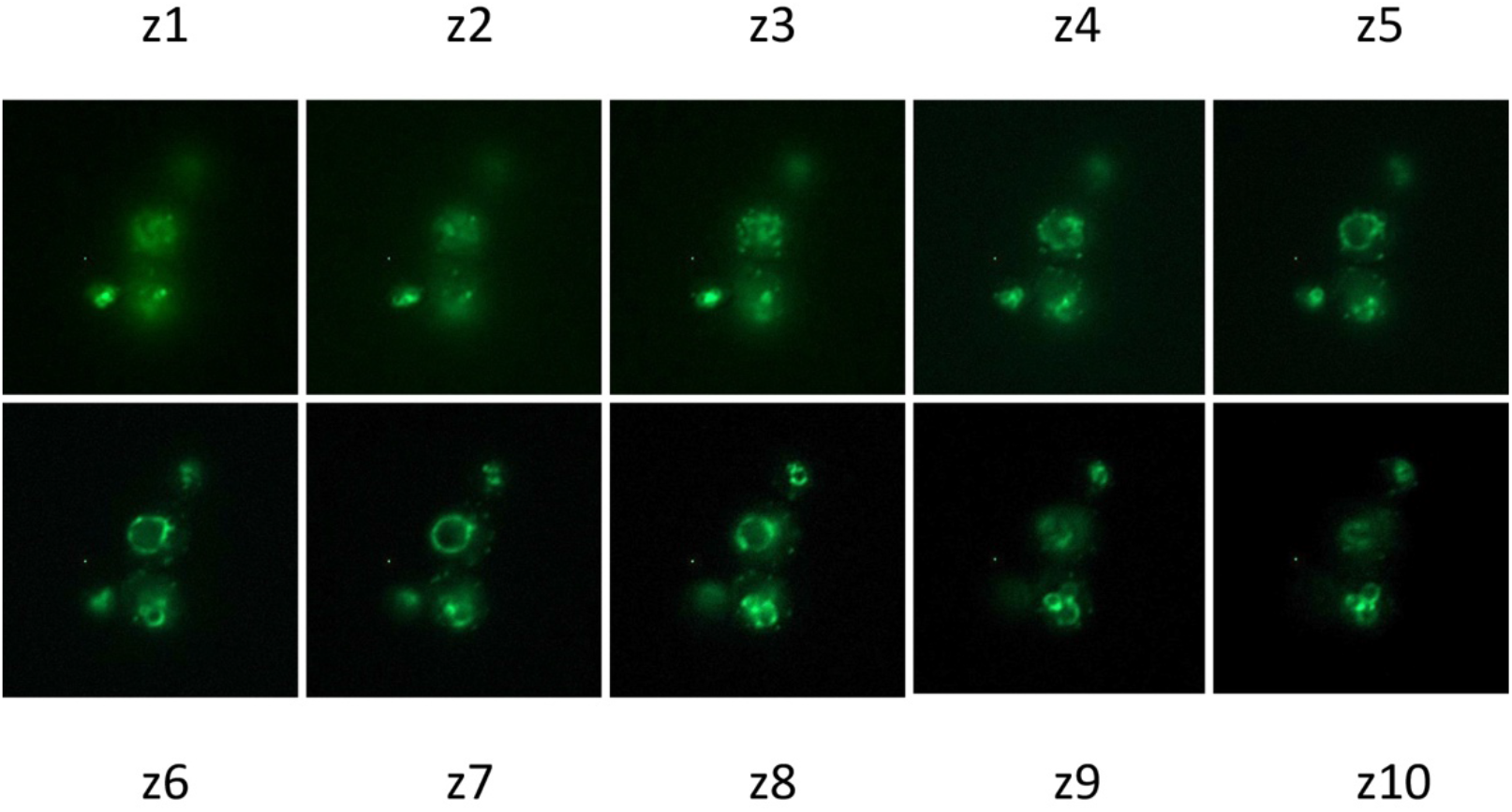
Z-stack of Mos10-sfGFP staining. Ten slices through RKY3216 cells expressing Mos10-sfGFP (z1 – z10).

**Figure S8-1.**
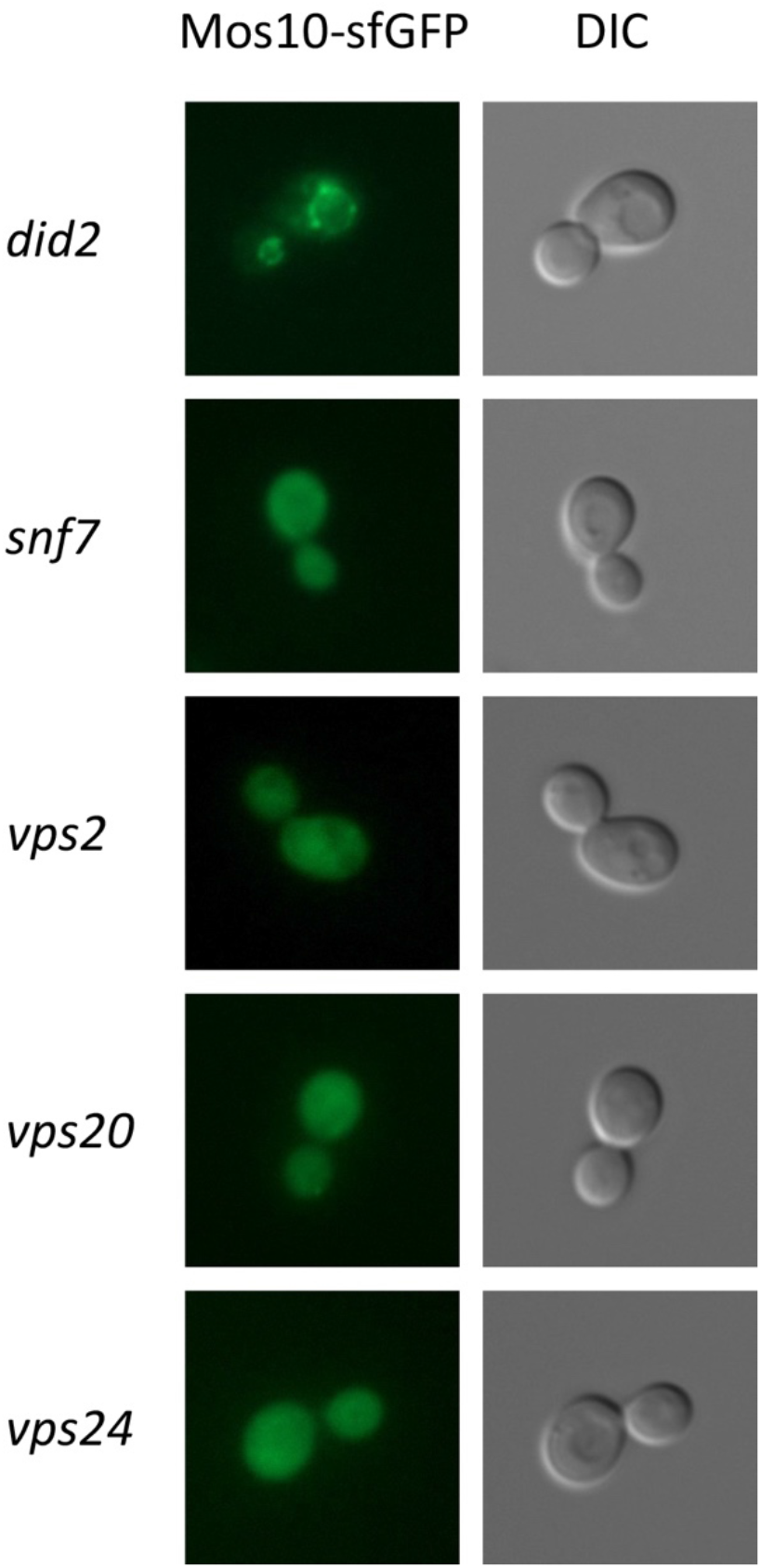
Effect of ESCRT-III deletions on the localization of Mos10-sfGFP. Several ESCRT-III genes were deleted in a strain expressing Mos10-sfGFP from its chromosomal locus. Left panel: Mos10-sfGFP fluorescence, right panel: DIC image. Strains from top to bottom (all expressing Mos10-sfGFP): RKY3478 (*Δdid2*), RKY3458 (*Δsnf7*), RKY3479 (*Δvps2*), RKY3437 (*Δvps20*), RKY3490 (*Δvps24*).

**Figure S8-2.**
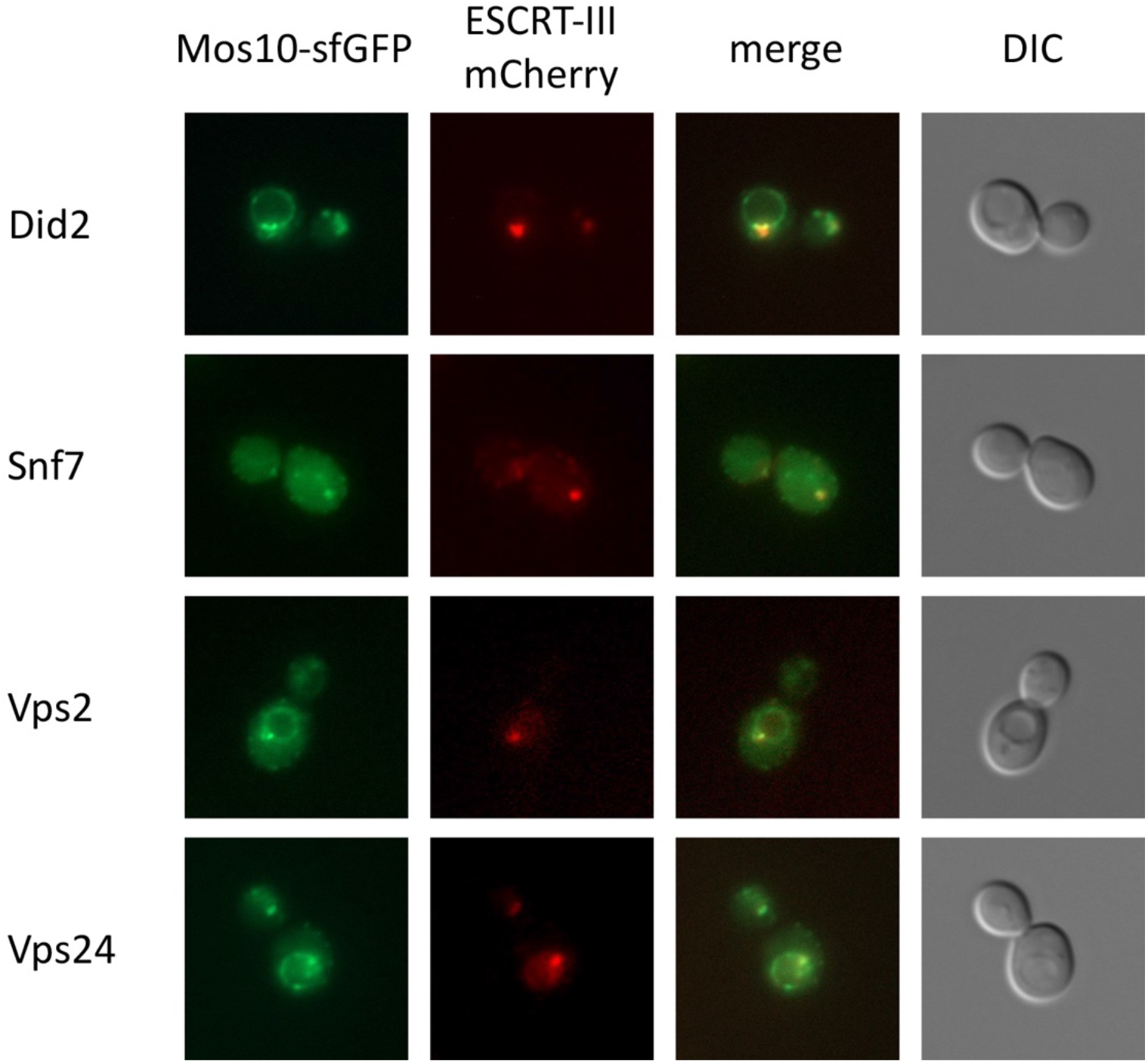
Co-localization of Mos10-sfGFP with ESCRT-III-mCherry fusions. Yeast strains expressing Mos10-sfGFP and ESCRT-III-mCherry fusions from their chromosomal loci were examined for co-localization by fluorescence microscopy. Panels from left to right: Mos10-sfGFP fluorescence, ESCRT-III-mCherry fluorescence, merged image, DIC image. Strains from top to bottom (all expressing Mos10-sfGFP): RKY3481 (Did2-mCherry), RKY3459 (Snf7-mCherry), RKY3461 (Vps2-mCherry), RKY3462 (Vps24-mCherry).

**Tab. S1.** Mos10 interacting proteins identified by SILAC/MS (n=3)

| Protein | CRAPome* | H/L ratio |  |  |
| --- | --- | --- | --- | --- |
| **VPS60** | **0 / 17** | **115.8** | **13.7** | **13.0** |
| TPS1 | 0 / 17 | 22.4 |  |  |
| COG8 | 1 / 17 | 11.1 |  |  |
| MSS1 | 0 / 17 | 11.0 |  |  |
| **VPS2** | **0 / 17** | **10.6** | **10.5** | **8.2** |
| *RPS3* | *17 / 17* | *9.7* |  |  |
| TAF7 | 0 / 17 | 9.2 |  |  |
| **SNF7** | **0 / 17** | **8.8** | **8.2** | **7.4** |
| PRE6 | 2 / 17 | 7.8 |  |  |
| **VPS24** | **0 / 17** | **7.7** | **4.0** | **2.6** |
| NTH1 | 0 / 17 | 7.7 |  |  |
| EXO84 | 1 / 17 | 6.5 |  |  |
| *FKS1* | *7 / 17* | *5.7* |  |  |
| *SSB1* | *17 / 17* | *5.7* |  |  |
| VMA5 | 4 / 17 | 5.5 |  |  |
| **VPS68** | **0 / 17** | **4.8** |  |  |
| **DID2** | **0 / 17** | **4.5** | **4.2** |  |
| ARP3 | 1 / 17 | 4.5 |  |  |
| DNF2 | 1 / 17 | 4.1 |  |  |
| *HEF3* | *8 / 17* | *3.9* |  |  |
| *NEW1* | *11 / 17* | *3.6* |  |  |
| YKR075C<sup>+</sup> | 0 / 17 | 3.6 |  |  |
| YGR130C | 0 / 17 | 3.3 |  |  |
| *RPL9B, RPL9A* | *17 / 17* | *3.2* |  |  |
| RTS1 | 0 / 17 | 3.2 |  |  |
| YCK2 | 1 / 17 | 3.1 |  |  |
| OAC1 | 0 / 17 | 2.9 | 2.3 |  |
| CBK1 | 4 / 17 | 2.9 |  |  |
| PTR2 | 0 / 17 | 2.9 |  |  |
| ALD5 | 1 / 17 | 2.8 |  |  |
| LEU1 | 1 / 17 | 2.7 |  |  |
| HBT1 | 0 / 17 | 2.5 | 2.3 |  |
| *HHT1* | *11 / 17* | *2.5* |  |  |
| MNN5 | 0 / 17 | 2.5 |  |  |
| PAN1 | 1 / 17 | 2.5 |  |  |
| *PMA1, PMA2* | *17 / 17* | *2.5* |  |  |
| FRT2 | 0 / 17 | 2.4 |  |  |
| *HXK1, HXK2* | *6 / 17* | *2.4* |  |  |
| YTA12 | 0 / 17 | 2.4 |  |  |
| BAG7<sup>+</sup> | 0 / 17 | 2.3 |  |  |
| CDC28 | 0 / 17 | 2.3 |  |  |
| STB6 | 0 / 17 | 2.3 |  |  |
| HXT1 | 0 / 17 | 2.2 |  |  |
| *TDH1* | *12 / 17* | *2.2* |  |  |
| TIM10 | 1 / 17 | 2.2 |  |  |
| TYA | n.a. | 2.2 |  |  |
| COX4 | 0 / 17 | 2.1 |  |  |
| TOM71 | 0 / 17 | 2.1 |  |  |
| YCK1 | 0 / 17 | 2.1 |  |  |
| *ILV5* | *6 / 17* | *2.0* |  |  |
| *SSE2* | *5 / 17* | *2.0* |  |  |
| YAT2 | 0 / 17 | 2.0 |  |  |

**ESCRT-III proteins and Vps68 in bold print**
\*The CRAPome: a Contaminant Repository for Affinity Purification Mass Spectrometry Data. Mellacheruvu et al. (2013) Nature Methods 10, 730–736
CRAPome > 4/17 (detected 4-times or more in 17 purifications) highlighted
\*Poly-His Proteins

**Tab S2:** Yeast strains

| Strain | Genotype |  |
| --- | --- | --- |
| JD52 | *MATa ura3-52 his3-Δ200 leu2-3,112 trp1-Δ63 lys2-801 ssd1-d2* | J. Dohmen,<br>Köln |
| STY50 | *MATα his4-401 leu2-3,112 trp1-1 ura3-52 HOL1-1* | R. Schneiter,<br>Fribourg |
| RKY 1920 | *MATa Δvps21::LEU2* | this study |
| RKY 2790 | *MATa Δsnf7* | this study |
| RKY 2830 | *MATa Δvps24* | this study |
| RKY 2889 | *MATa MOS10-6His::His3* | this study |
| RKY 2909 | *MATa Δmos10::His3* | this study |
| RKY 2978 | *MATa SNF7-6His::His3* | this study |
| RKY 2998 | *MATa Δarg4* | this study |
| RKY 2999 | *MATa Δarg4 MOS10-6His::His3* | this study |
| RKY 3183 | *MATa VPS68-sfGFP::kan* | this study |
| RKY 3210 | *MATa Δarg4 MOS10-sfGFP::kan* | this study |
| RKY 3214 | *MATa DID2-sfGFP::kan* | this study |
| RKY 3215 | *MATa IST1-sfGFP::kan* | this study |
| RKY 3216 | *MATa MOS10-sfGFP::kan* | this study |
| RKY 3217 | *MATa SNF7-sfGFP::kan* | this study |
| RKY 3218 | *MATa VPS2-sfGFP::kan* | this study |
| RKY 3220 | *MATa VPS24-sfGFP::kan* | this study |
| RKY 3222 | *MATa Δvps68::kan* | this study |
| RKY 3224 | *MATa MOS10-sfGFP::kan Δvps68::His3* | this study |
| RKY 3257 | *MATa IST1-sfGFP::His3 Δvps68::kan* | this study |
| RKY 3269 | *MATa DID2-sfGFP::kan Δvps68::His3* | this study |
| RKY 3271 | *MATa VPS2-sfGFP::kan Δvps68::His3* | this study |
| RKY 3274 | *MATa VPS24-sfGFP::kan Δvps68::His3* | this study |
| RKY 3279 | *MATa SNF7-sfGFP::kan Δvps68::His3* | this study |
| RKY 3285 | *MATa vps68::[kan SNF7p-sfGFP-VPS68]* | this study |
| RKY 3300 | *MATa Δsnf7 Δvps68::kan* | this study |
| RKY 3307 | *MATa Δvps21::LEU2 Δvps68::kan* | this study |
| RKY 3319 | *MATa ste6::[kan GAL1p-STE6]* | this study |
| RKY 3320 | *MATa Δvps68::kan ste6::[His3 GAL1p-STE6]* | this study |
| RKY 3351 | *MATa ypt10::[His3 SNF7p-sfGFP-YPT10]* | this study |
| RKY 3361 | *MATa SNF7-mCherry::kan ypt53::[His3 SNF7p-sfGFP-YPT53]* | this study |
| RKY 3362 | *MATa SNF7-mCherry::kan ypt31::[His3 SNF7p-sfGFP-YPT31]* | this study |
| RKY 3363 | *MATa SNF7-mCherry::kan ypt32::[His3 SNF7p-sfGFP-YPT32]* | this study |
| RKY 3364 | *MATa SNF7-mCherry::kan ypt6::[His3 SNF7p-sfGFP-YPT6]* | this study |
| RKY 3365 | *MATa SNF7-mCherry::kan ypt10::[His3 SNF7p-sfGFP-YPT10]* | this study |
| RKY 3376 | *MATa Δvps21::LEU2 Δsnf7::kan* | this study |
| RKY 3384 | *MATa DID2-mCherry::kan SNF7-sfGFP::His* | this study |
| RKY 3394 | *MATa vps68::[kan SNF7p-sfGFP-VPS68] Δmos10::His3* | this study |
| RKY 3412 | *MATa vps68::[kan SNF7p-sfGFP-VPS68] Δsnf7::His3* | this study |
| RKY 3423 | *MATa SNF7-mCherry::kan vps21::[His3 SNF7p-sfGFP-VPS21]* | this study |
| RKY 3425 | *MATa SNF7-mCherry::kan ypt52::[His3 SNF7p-sfGFP-YPT52]* | this study |
| RKY 3429 | *MATa MOS10-sfGFP::kan SEC63-mCherry::His3* | this study |
| RKY 3437 | *MATa MOS10-sfGFP::kan Δvps20::His3* | this study |
| RKY 3438 | *MATa MOS10-6His::His3 Δsnf7::kan* | this study |
| RKY 3448 | *MATa MOS10-sfGFP::kan VPH1-mCherry::His3* | this study |
| RKY 3458 | *MATa MOS10-sfGFP::kan Δsnf7::His3* | this study |
| RKY 3459 | *MATa MOS10-sfGFP::kan SNF7-mCherry::His3* | this study |
| RKY 3461 | *MATa MOS10-sfGFP::kan VPS2-mCherry::His3* | this study |
| RKY 3462 | *MATa MOS10-sfGFP::kan VPS24-mCherry::His3* | this study |
| RKY 3473 | *MATa MOS10-mCherry::kan vps21::[His3 SNF7p-sfGFP-VPS21]* | this study |
| RKY 3478 | *MATa MOS10-sfGFP::kan Δdid2::His3* | this study |
| RKY 3479 | *MATa MOS10-sfGFP::kan Δvps2::His3* | this study |
| RKY 3481 | *MATa MOS10-sfGFP::His3 DID2-mCherry::kan* | this study |
| RKY 3482 | *MATa MOS10-mCherry::His3 COG5-sfGFP::kan* | this study |
| RKY 3484 | *MATa MOS10-mCherry::His3 cps1::[kan SNF7p-sfGFP-CPS1]* | this study |
| RKY 3486 | *MATa MOS10-mCherry::His3 STE6-sfGFP::kan* | this study |
| RKY 3490 | *MATa MOS10-sfGFP::kan Δvps24* | this study |

**Tab. S3.** Plasmids

| Plasmid | Insert | Vector* | Replicon, Marker |
| --- | --- | --- | --- |
| pRK599 | STE6-GFP | YEplac195 | 2μ, URA3 |
| pRK1408 | HXT7p-mCherry-CPS1 | YCplac22 | CEN, TRP1 |
| pRK2015 | PDC1p-3HA-SUC2-HIS4C | YCplac33 | CEN, URA3 |
| pRK2016 | PDC1p-VPS68 (TM1)-3HA-SUC2-HIS4C | YCplac33 | CEN, URA3 |
| pRK2017 | PDC1p-VPS68 (TM1-2)-3HA-SUC2-HIS4C | YCplac33 | CEN, URA3 |
| pRK2018 | PDC1p-VPS68 (TM1-3)-3HA-SUC2-HIS4C | YCplac33 | CEN, URA3 |
| pRK2019 | PDC1p-VPS68 (TM1-4)-3HA-SUC2-HIS4C | YCplac33 | CEN, URA3 |

\* Gietz and Sugino, 1988

## References

Adell MAY, Migliano SM, Upadhyayula S, Bykov YS, Sprenger S, Pakdel M, Vogel GF, Jih G, Skillern W, Behrouzi R et al (2017) Recruitment dynamics of ESCRT-III and Vps4 to endosomes and implications for reverse membrane budding. Elife 6

Azmi IF, Davies BA, Xiao J, Babst M, Xu Z, Katzmann DJ (2008) ESCRT-III family members stimulate Vps4 ATPase activity directly or via Vta1. Dev Cell 14: 50–61

Babst M, Katzmann DJ, Estepa-Sabal EJ, Meerloo T, Emr SD (2002) ESCRT-III: an endosome-associated heterooligomeric protein complex required for MVB sorting. Dev Cell 3: 271–282

Banjade S, Shah YH, Tang S, Emr SD (2021) Design principles of the ESCRT-III Vps24-Vps2 module. Elife 10

Banjade S, Tang S, Shah YH, Emr SD (2019) Electrostatic lateral interactions drive ESCRT-III heteropolymer assembly. Elife 8

Bauer I, Brune T, Preiss R, Kölling R (2015) Evidence for a non-endosomal function of the *Saccharomyces cerevisiae* ESCRT-III like protein Chm7. Genetics 201: 1439–1452

Bertin A, de Franceschi N, de la Mora E, Maity S, Alqabandi M, Miguet N, di Cicco A, Roos WH, Mangenot S, Weissenhorn W et al (2020) Human ESCRT-III polymers assemble on positively curved membranes and induce helical membrane tube formation. Nat Commun 11: 2663

Bonangelino CJ, Chavez EM, Bonifacino JS (2002) Genomic screen for vacuolar protein sorting genes in Saccharomyces cerevisiae. Mol Biol Cell 13: 2486–2501

Bowers K, Stevens TH (2005) Protein transport from the late Golgi to the vacuole in the yeast Saccharomyces cerevisiae. Biochim Biophys Acta 1744: 438–454

Brune T, Kunze-Schumacher H, Kölling R (2019) Interactions in the ESCRT-III network of the yeast Saccharomyces cerevisiae. Curr Genet 65: 607–619

Chiaruttini N, Redondo-Morata L, Colom A, Humbert F, Lenz M, Scheuring S, Roux A (2015) Relaxation of Loaded ESCRT-III Spiral Springs Drives Membrane Deformation. Cell 163: 866–879

Dimaano C, Jones CB, Hanono A, Curtiss M, Babst M (2008) Ist1 regulates Vps4 localization and assembly. Mol Biol Cell 19: 465–474

Gatta AT, Carlton JG (2019) The ESCRT-machinery: closing holes and expanding roles. Curr Opin Cell Biol 59: 121–132

Gautier R, Douguet D, Antonny B, Drin G (2008) HELIQUEST: a web server to screen sequences with specific alpha-helical properties. Bioinformatics 24: 2101–2102

Gimenez-Andres M, Copic A, Antonny B (2018) The Many Faces of Amphipathic Helices. Biomolecules 8

Heinzle C, Mücke L, Brune T, Kölling R (2019) Comprehensive analysis of yeast ESCRT-III composition in single ESCRT-III deletion mutants. Biochem J 476: 2031–2046

Huh WK, Falvo JV, Gerke LC, Carroll AS, Howson RW, Weissman JS, O’Shea EK (2003) Global analysis of protein localization in budding yeast. Nature 425: 686–691

Hurley JH (2015) ESCRTs are everywhere. EMBO J 34: 2398–2407

Junglas B, Huber ST, Heidler T, Schlosser L, Mann D, Hennig R, Clarke M, Hellmann N, Schneider D, Sachse C (2021) PspA adopts an ESCRT-III-like fold and remodels bacterial membranes. Cell 184: 3674–3688 e3618

Kölling R, Hollenberg CP (1994) The ABC-transporter Ste6 accumulates in the plasma membrane in a ubiquitinated form in endocytosis mutants. EMBO J 13: 3261–3271

Kranz A, Kinner A, Kölling R (2001) A family of small coiled-coil-forming proteins functioning at the late endosome in yeast. Mol Biol Cell 12: 711–723

Krsmanović T, Pavelec A, Sydor T, Kölling R (2005) Control of Ste6 recycling in the early endocytic pathway in yeast. Mol Biol Cell 16: 2809–2821

Langemeyer L, Borchers AC, Herrmann E, Fullbrunn N, Han Y, Perz A, Auffarth K, Kummel D, Ungermann C (2020) A conserved and regulated mechanism drives endosomal Rab transition. Elife 9

Lee IH, Kai H, Carlson LA, Groves JT, Hurley JH (2015) Negative membrane curvature catalyzes nucleation of endosomal sorting complex required for transport (ESCRT)-III assembly. Proc Natl Acad Sci U S A 112: 15892–15897

Leung KF, Dacks JB, Field MC (2008) Evolution of the multivesicular body ESCRT machinery; retention across the eukaryotic lineage. Traffic 9: 1698–1716

Liu J, Tassinari M, Souza DP, Naskar S, Noel JK, Bohuszewicz O, Buck M, Williams TA, Baum B, Low HH (2021) Bacterial Vipp1 and PspA are members of the ancient ESCRT-III membrane-remodeling superfamily. Cell 184: 3660–3673 e3618

Loibl M, Grossmann G, Stradalova V, Klingl A, Rachel R, Tanner W, Malinsky J, Opekarova M (2010) C terminus of Nce102 determines the structure and function of microdomains in the Saccharomyces cerevisiae plasma membrane. Eukaryot Cell 9: 1184–1192

Longtine MS, McKenzie A, Demarini DJ, Shah NG, Wach A, Brachat A, Philippsen P, Pringle JR (1998) Additional modules for versatile and economical PCR-based gene deletion and modification in *Saccharomyces cerevisiae*. Yeast 14: 953–961

Maity S, Caillat C, Miguet N, Sulbaran G, Effantin G, Schoehn G, Roos WH, Weissenhorn W (2019) VPS4 triggers constriction and cleavage of ESCRT-III helical filaments. Sci Adv 5: eaau7198

McCullough J, Frost A, Sundquist WI (2018) Structures, Functions, and Dynamics of ESCRT-III/Vps4 Membrane Remodeling and Fission Complexes. Annu Rev Cell Dev Biol 34: 85–109

Mellacheruvu D, Wright Z, Couzens AL, Lambert JP, St-Denis NA, Li T, Miteva YV, Hauri S, Sardiu ME, Low TY et al (2013) The CRAPome: a contaminant repository for affinity purification-mass spectrometry data. Nat Methods 10: 730–736

Mierzwa BE, Chiaruttini N, Redondo-Morata L, von Filseck JM, Konig J, Larios J, Poser I, Muller-Reichert T, Scheuring S, Roux A et al (2017) Dynamic subunit turnover in ESCRT-III assemblies is regulated by Vps4 to mediate membrane remodelling during cytokinesis. Nat Cell Biol 19: 787–798

Nickerson DP, West M, Henry R, Odorizzi G (2010) Regulators of Vps4 ATPase activity at endosomes differentially influence the size and rate of formation of intralumenal vesicles. Mol Biol Cell 21: 1023–1032

Nickerson DP, West M, Odorizzi G (2006) Did2 coordinates Vps4-mediated dissociation of ESCRT-III from endosomes. J Cell Biol 175: 715–720

Pawelec A, Arsić J, Kölling R (2010) Mapping of Vps21 and HOPS binding sites in Vps8 and effect of binding site mutants on endocytic trafficking. Eukaryot Cell 9: 602–610

Pfitzner AK, Mercier V, Jiang X, Moser von Filseck J, Baum B, Saric A, Roux A (2020) An ESCRT-III Polymerization Sequence Drives Membrane Deformation and Fission. Cell 182: 1140–1155 e1118

Raymond CK, Howald SI, Vater CA, Stevens TH (1992) Morphological classification of the yeast vacuolar protein sorting mutants: evidence for a prevacuolar compartment in class E *vps* mutants. Mol Biol Cell 3: 1389–1402

Rue SM, Mattei S, Saksena S, Emr SD (2008) Novel Ist1-Did2 complex functions at a late step in multivesicular body sorting. Mol Biol Cell 19: 475–484

Schluter C, Lam KK, Brumm J, Wu BW, Saunders M, Stevens TH, Bryan J, Conibear E (2008) Global analysis of yeast endosomal transport identifies the vps55/68 sorting complex. Mol Biol Cell 19: 1282–1294

Sengstag C, Stirling C, Schekman R, Rine J (1990) Genetic and biochemical evaluation of eucaryotic membrane protein topology: multiple transmembrane domains of Saccharomyces cerevisiae 3-hydroxy-3-methylglutaryl coenzyme A reductase. Mol Cell Biol 10: 672–680

Tang S, Henne WM, Borbat PP, Buchkovich NJ, Freed JH, Mao Y, Fromme JC, Emr SD (2015) Structural basis for activation, assembly and membrane binding of ESCRT-III Snf7 filaments. Elife 4

Teis D, Saksena S, Emr SD (2008) Ordered assembly of the ESCRT-III complex on endosomes is required to sequester cargo during MVB formation. Dev Cell 15: 578–589

Thaller DJ, Allegretti M, Borah S, Ronchi P, Beck M, Lusk CP (2019) An ESCRT-LEM protein surveillance system is poised to directly monitor the nuclear envelope and nuclear transport system. Elife 8

Vietri M, Radulovic M, Stenmark H (2020) The many functions of ESCRTs. Nat Rev Mol Cell Biol 21: 25–42

